# Altered astrocytic and microglial homeostasis characterizes a decreased proinflammatory state in bipolar disorder

**DOI:** 10.1101/2023.10.29.564621

**Authors:** Quentin Amossé, Benjamin B. Tournier, Aurélien M. Badina, Lilou Marchand-Maillet, Laurene Abjean, Sylvain Lengacher, Nurun Fancy, Amy M. Smith, Yeung-Yeung Leung, Verena Santer, Valentina Garibotto, David R. Owen, Camille Piguet, Kelly Ceyzériat, Stergios Tsartsalis, Philippe Millet

**Author notes:** These authors share senior authorship (Stergios Tsartsalis, Philippe Millet). Address for correspondence: Dr Stergios Tsartsalis Department of Psychiatry University Hospitals of Geneva Chemin du Petit-Bel-Air 2 CH1226, Thônex, Switzerland; Prof Philippe Millet Department of Psychiatry University Hospitals of Geneva, Fondation pour recherches medicales Avenue de la Roseraie 64, CH1205, Geneva, Switzerland.

## Abstract

Multiple lines of evidence point to peripheral immune alterations in bipolar disorder (BD) although the activity of brain immune mechanisms remain largely unexplored. To identify the cell type-specific immune alterations in the BD brain, we performed a proteomic and single nuclear transcriptomic analysis of *postmortem* cingulate cortex samples from BD and control subjects. Our results showed that genes associated to the genetic risk for BD are enriched in microglia and astrocytes. Transcriptomic alterations in microglia point to a reduced proinflammatory phenotype, associated to reduced resistance to oxidative stress and apoptosis, which was confirmed with immunohistochemical quantification of IBA1 density. Astrocytes show transcriptomic evidence of an imbalance of multiple metabolic pathways, extracellular matrix composition and downregulated immune signalling. These alterations are associated to *ADCY2* and *NCAN,* two GWAS genes upregulated in astrocytes. Finally, cell-cell communication analysis prioritized upregulated SPP1-CD44 signalling to astrocytes as a potential regulator of the transcriptomic alterations in BD. Our results indicate that microglia and astrocytes are characterized by downregulated immune responses associated to a dysfunction of core mechanisms via which these cells contribute to brain homeostasis.

## Introduction

Bipolar disorder (BD) is a severe and highly heritable psychiatric disorder, with significant morbidity^1^, affecting 1% to 4.5% of the population^2^ and is characterized by episodes of mood alteration, that can be depressive or (hypo)manic. A significant proportion of BD patients suffers from functional and cognitive impairment at various degrees across the phases of the disease^3^.

The aetiology of the disease is still largely unknown, but it is considered multifactorial^4^, encompassing genetic and environmental factors^5,6^. The absence of reliable animal models of the disease (based on genetic or environmental mechanisms) further hampers the understanding of the pathology. Indeed, the clinical heterogeneity of BD patients combined with the high comorbidity rate leads to difficulties in the identification of specific biological targets. Nevertheless, evidence from Genome Wide Association Studies (GWAS)^7–10^ have highlighted several genomic loci with a significant association with BD.

One of the potential biological mechanisms that has been widely described in BD is inflammation. Indeed, an increased concentration of peripheral inflammatory markers has been consistently reported in cross-sectional studies in BD, suggesting immune alterations across the various mood states of the disease^1,11–13^. However, whether this increased peripheral inflammatory state is due to central inflammation has not been determined. If peripheral inflammation is mechanistically associated with BD pathophysiology central immune mechanisms should also be activated. This central immune activation should involve microglia and astrocytes, which are among the most important cellular mediators of immunity in the central nervous system (CNS)^14^. Contrary to the upregulation of peripheral inflammatory markers in BD, a recent large-scale bulk RNA sequencing study of over 500 brain samples from two brain regions involved in BD pathophysiology (cingulate cortex and amygdala) has shown that gene co-expression modules enriched in immune pathways were significantly downregulated in BD^15^. However, bulk RNA sequencing does not allow for cell type-specific inferences to be made, so it is impossible to assert if this downregulation of immune pathways involves microglia and astrocytes. Furthermore, a recent systematic review of human neuropathology studies assessing immune markers in *postmortem* human brain samples highlighted that at the protein level, results are inconsistent across studies^16^, likely due to the heterogeneity of the employed techniques and the targeted molecules.

To clarify the central molecular alterations involved in BD, we first performed a bulk tissue proteomic analysis in human *postmortem* samples from BD patients and age- and sex-matched controls, which showed that proteomic alterations in BD involved astrocyte- and microglia-specific molecules. Consequently, using single nucleus RNA sequencing (snRNAseq), we focused on microglia and astrocytes and first assessed the astrocyte- and microglia-specific expression of BD GWAS genes and their alterations with the disease. We then characterized the wider dysfunctional biological pathways in BD using snRNAseq and spatial transcriptomics and validated the most important findings using immunohistology and an independent bulk tissue transcriptomic dataset. Our multi-omic analyses results point to specific molecular alterations in glial cells in BD, predominantly characterized by extracellular matrix (ECM) gene overexpression, metabolic and immune signalling dysfunction in astrocytes and reduced proinflammatory, phagocytosis-related signalling and apoptosis-associated decrease in microglial density in BD.

## Results

### BD is characterized by proteomic alteration of immune, mitochondrial and lipid metabolic pathways

To identify protein pathway alterations in BD, we first performed a proteomic analysis on bulk cingulate gyrus (Cg) tissue–a brain region that has been consistently implicated in BD – from 18 *postmortem* human brain samples from BD patients and age- and sex-matched non-neurological and non-psychiatric control subjects (CT). To increase the statistical power in the assessment of protein functional pathways that are altered in BD, we employed a protein co-expression analysis using weighted gene co-expression network analysis (WGCNA)^17^, which yielded 13 modules. WGCNA uses principal component analysis (PCA) to extract an eigenprotein value, i.e. a numeric value representing the overall abundance of all the proteins in the module (Table S1). We thus compared the eigenprotein values of the modules between the BD and CT samples and based on this comparison we prioritized the two significantly differentially abundant modules: Module 17 (M17, 104 proteins, *p_adj_* = 0.0194, downregulated in BD, Figure 1A) and, “Module” 0 (M0, 185 proteins, *p_adj_* = 0.0184, upregulated in BD), which actually is the set of proteins that were not assigned to any module in the WGCNA analysis (Figure 1B). Functional enrichment analysis (FEA) of the M17 module using enrichR^18^ showed enrichment in growth factor pathways (involving proteins that mediate VEGF, NTRK2-NTRK3, FGF and PDGFB signalling), innate immune/phagocytosis pathways and citrate cycle proteins and mitochondrial matrix proteins (Figure 1C). The presence of C1S in M17 indicates a downregulation of the classical pathway of complement activation, whereas the presence of the complement regulator CD55 indicates a potential mechanism for neurotoxicity in BD^19^ and corroborates previous findings of a downregulation of CD55 in the plasma of BD patients^20^. We then performed individual protein level differential abundance analysis focusing on proteins included in this module to prioritize potential drivers of its downregulation. ATP-dependent 6-phosphofructokinase (PFKL), isocitrate dehydrogenase (IDH3G), mitochondrial acetyl-coenzyme A synthetase 2-like (ACSS1) and mitochondrial ornithine aminotransferase (OAT) were significantly downregulated (log_2_FC_PKFL_ = −0.313, *p_adjPKFL_* < 0.0001, log_2_FC_ACSS1_ = −0.303, *p_adjACSS1_* = 0.00481, log_2_FC_OAT_ = −0.251, *p_adjOAT_* = 0.0164), underlining the dysregulation of mitochondrial and metabolic function in BD^21,22^ (Figure 1D). In the M0 protein set, lipoprotein metabolism, ECM, apoptosis and neurotransmitter signalling pathways were enriched (Figure 1E). Individual protein differential abundance analysis highlighted lipid processing alterations (APOE, APOA1 and FABP3) as significantly upregulated in BD (log_2_FC_APOE_ = 0.373, *p_adjAPOE_* < 0.0001, log_2_FC_APOA1_ = 0.431, *p_adjAPOA1_* = 0.0491, log_2_FC_FABP3_ = 0.333, *p_adjFABP3_* = 0.0293) (Figure 1D). Some of the donors of our BD group were taken psychotropic medications (lithium, olanzapine or valproate) at the time of death. We thus asked if the proteomic alterations in our samples could be driven by these medications. We identified gene sets that were in the brain of rodents in response to treatment with these medications from previous studies. We thus tested the overlap between the differentially abundant proteins in our samples with genes altered in response to lithium^23^, olanzapine^24^ or valproate^25^ and did not find any significant overlap. This results suggests that the results of our proteomic analysis is not driven by medications.

**Figure 1.**
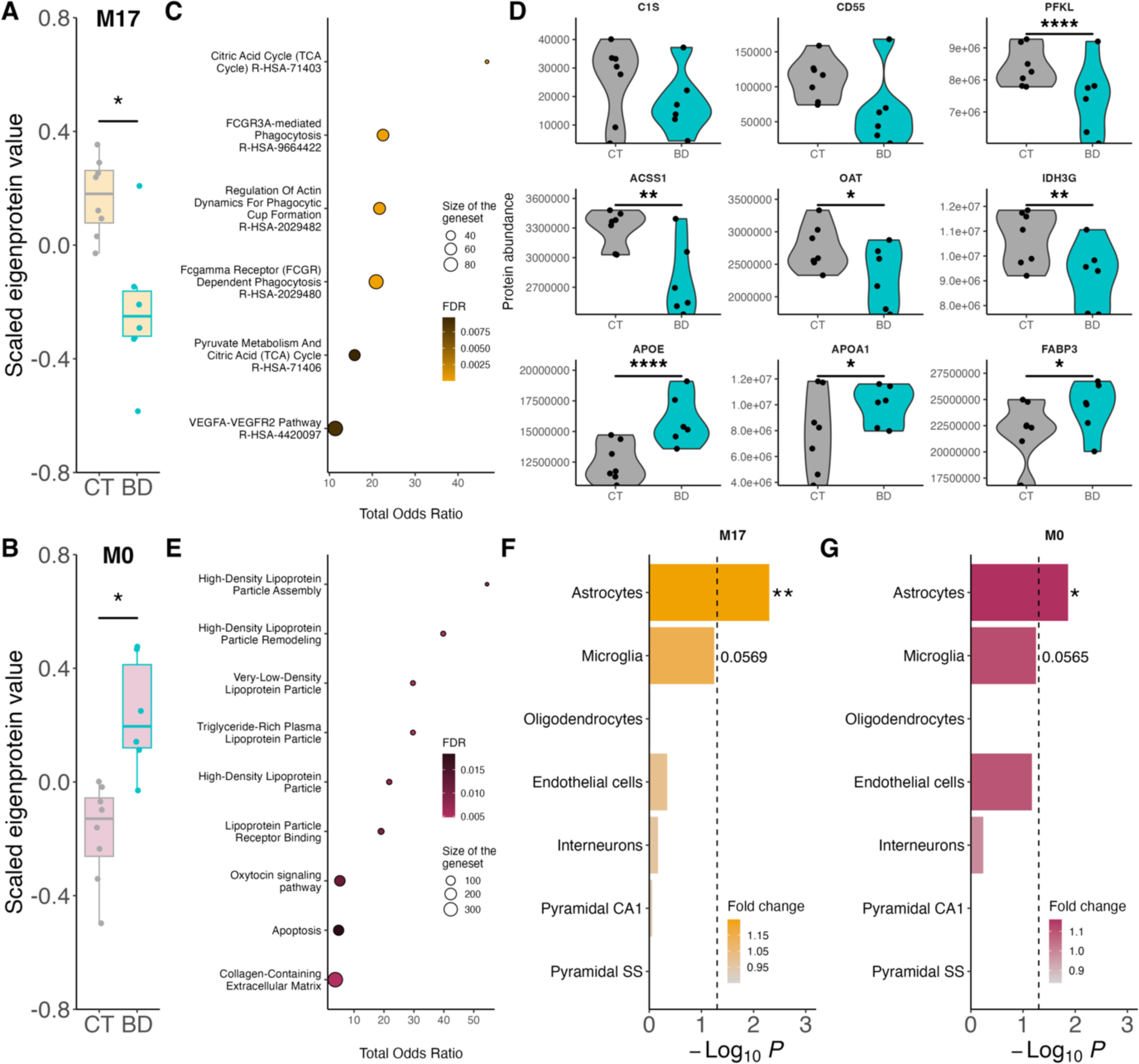
Proteomic modules altered in BD are enriched in astrocytic and microglial genes (Cg). (**A-B**) Eigenprotein score of module 17 (M17, **A**) and module 0 (M0, **B**) proteins in control (CT) and bipolar disorder (BD) subjects in cingulate gyrus (Cg). (**C**) Functional enrichment analysis (FEA) results of the 104 proteins of the M17. (**D**) Differential abundance of individual proteins in bulk proteomics PFKL (p_adj_ < 0.0001, LogFC = −0.313), ACSS1 (p_adj_ = 0.00481, LogFC = −0.303), OAT (p_adj_ = 0.0164, LogFC = −0.251), IDH3G (p_adj_ = 0.00208, LogFC = −0.230), APOE (p_adj_ < 0.0001, LogFC = 0.373), APOA1 (p_adj_ = 0.0491, LogFC = 0.431), FABP3 (p_adj_ = 0.0293, LogFC = 0.333), (**E**) FEA results of the 185 proteins of the M0. (**F-G**) Expression Weighted Cell Type Enrichment (EWCE) results of the 104 genes composing the M17 (**F**) and of the 185 proteins composing the M0 (**G**).

Given that this proteomic analysis was performed on bulk tissue, we then sought to identify cell type associations of the significantly altered protein sets. We employed the Expression Weighted Cell type Enrichment (EWCE) method^26^ that assesses if a given set of proteins/genes is significantly enriched in markers of a particular cell type based on brain single cell RNA sequencing datasets. This analysis identified a significant enrichment of astrocyte-specific proteins in both downregulated (M17) and upregulated (M0) sets of proteins (*p*_downregulated_ = 0.005, *p*_upregulated_ = 0.0137, respectively, Figure 1G, H). In both modules, microglia-specific enrichment approached significance in this test (*p*_downregulated_ = 0.0569, *p*_upregulated_ = 0.0565, Figure 1F, G). No other cell types were enriched in any of the modules. Taken together, these results suggest immune, lipid and mitochondrial metabolic alterations in BD that are specifically associated with astrocytes and microglia.

### BD risk factor genes are differentially expressed in astrocytes and microglia

Our proteomic analysis suggested that the major alterations in protein expression in BD occurred in astrocytes and microglia. We thus specifically examined these two cell types using snRNAseq on a subset of our samples. We used fluorescence activated cell sorting (FACS) on the isolated nuclei to remove SOX10-positive oligodendrocytes and NEUN-positive neurons^27,28^ prior to barcoding. With this method, 15’000 ‘double negative’ nuclei were retained per sample (=0.079 ±0.047% of total events – Figure S1A). The nuclei were then processed on the 10X Chromium controller and sequenced. Then, we used the nf-core/scflow pipeline to perform integration using Liger^29^, clustering with UMAP^30^, and cell identification, resulting in 18’176 nuclei in total (Figure 2A). BD and CT as well as male and female donor nuclei were well-mixed after integration (Figure S1B, C). The number of nuclei did not significantly differ between BD and CT (*p* = 0.1033, Figure S1D), or between male and female donors (*p* = 0.3415, Figure S1E). Nuclei sorting and automated cell type annotation was verified with canonical microglial and astrocytic cell markers and with enrichment of the nuclei for gene sets characteristic of human microglia^31^ and astrocytes^32^ using AUCell^33^, which confirmed the cell type identity (Figure 2B-D, Figure S1F, G).

**Figure 2.**
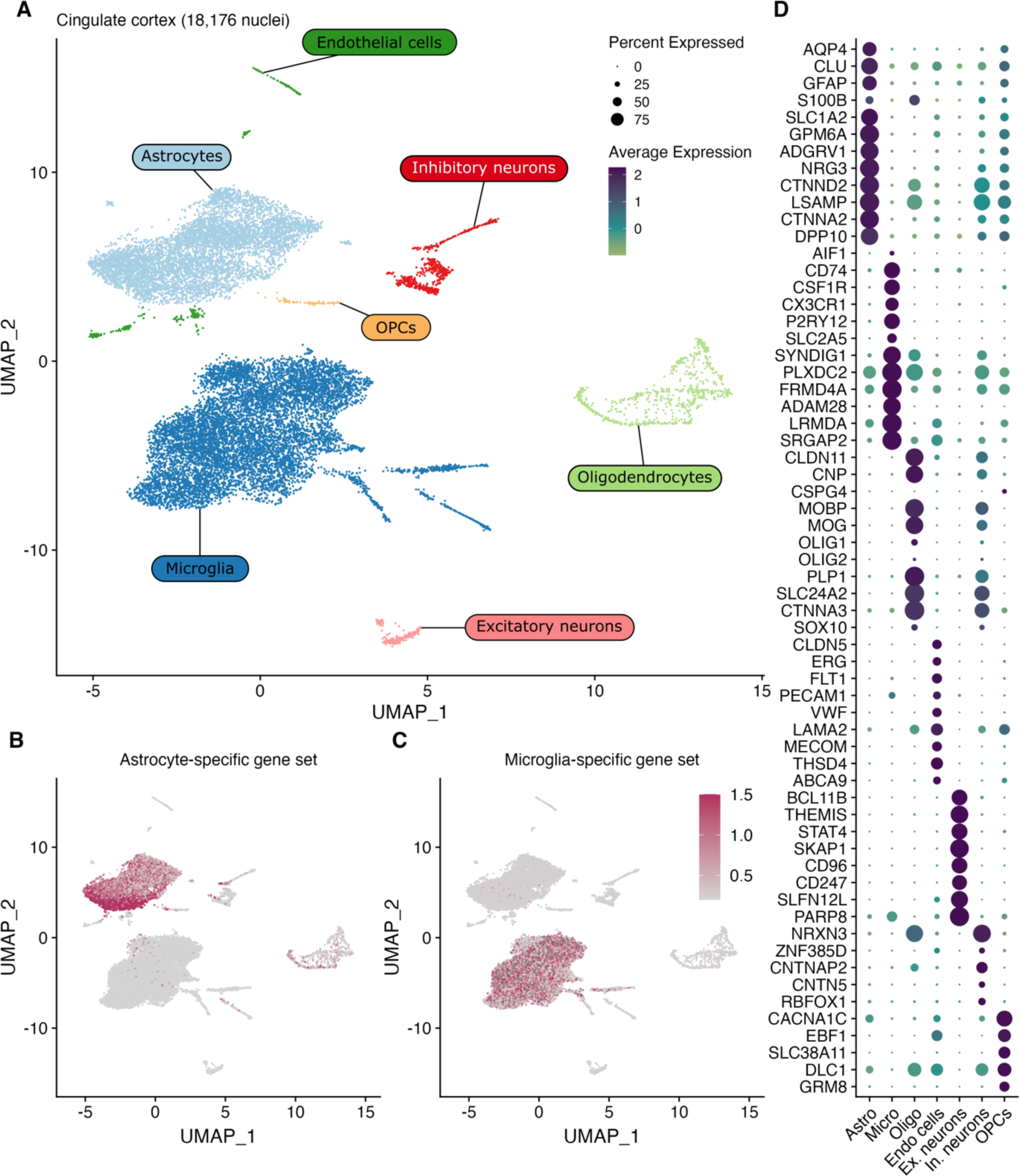
snRNAseq assessment of astrocytes and microglia. (**A**) UMAP 2D visualization of the 18,176 clustered nuclei from the cingulate cortex (average 1652 nuclei per sample), made up of 9868 (54.29%) microglia, 6386 (35.13%) astrocytes, 716 (3.94%) oligodendrocytes, 575 (3.16%) inhibitory neurons, 304 (1.67%) endothelial cells, 238 (1.31%) excitatory neurons and 89 (0.49%) OPCs. (**B**-**C**) Feature plot of the expression of the astrocyte-(**B**) and microglia-specific (**C**) gene sets, identifying the astrocytic and microglial clusters. (**D**) Dot plot representing the average expression of the various cluster-specific genes.

We first assessed if genes related to BD genetic risk-associated genomic loci from GWAS (a combined list of genes identified by Stahl et al.^7^ and Mullins et al.^10^, *Table S2*), were enriched in astrocytes and microglia nuclei. We first quantified the expression of the 73 GWAS genes. Of those genes, 43 are expressed by at least 5% of astrocytes or microglia in our study (Figure 3A). We then used MAST^34^ to perform a differential gene expression (DGE) analysis using a mixed effects model, between BD and CT samples and found that several of the GWAS genes were differentially expressed in glial cells (padj < 0.1, |logFC| > 0.1; Figure 3B, C): in astrocytes *ADCY2* (p_adj_= 0.092, logFC =0.698) coding for an isoform of adenylyl cyclase that catalyses the formation of cyclic adenosine monophosphate (cAMP) from ATP and is involved in cAMP-PKA signalling, regulating multiple cellular functions such as glycogen metabolism^35–38;^ *CACNB2* (p_adj_= 0.0669, logFC = −2.607), a subunit of a voltage-gated calcium channels which has been associated with hypertension^39,40^ and brain cognitive dysfunctions in BD^41,42^; *NCAN (p_adj_= 0.015, logFC = 0.785),* a component of brain ECM and specifically of brain perineuronal nets that regulate synaptic structure and function^43–47^, which also contributes to the formation of glial scar in response to injury^48–50;^ *SLC25A17* (p_adj_= 0.0369, logFC = 0.268), a peroxisomal transporter^51^; and *FSTL5* (p_adj_= 0.007, logFC = −0.352), coding for a Wnt signalling-associated protein^52^ shown to be implicated in caspase-dependent apoptosis^53^ (Figure 3B). In microglia, 3 genes were significantly altered: *SSBP2* (p_adj_= 0.00018, logFC = 0.754), a highly ubiquitously expressed gene coding for a single-stranded DNA binding protein^54^; *STK4* (p_adj_= 0.0231, logFC = −0.15), which is involved in microglial apoptosis and may contribute to microglial activation^55,56^; and *CD47* (p_adj_= 0.0856, logFC = −0.199), which codes for an inhibitor of microglial phagocytosis contributing to the regulation of synaptic pruning during development^57,58^ (Figure 3C).

**Figure 3.**
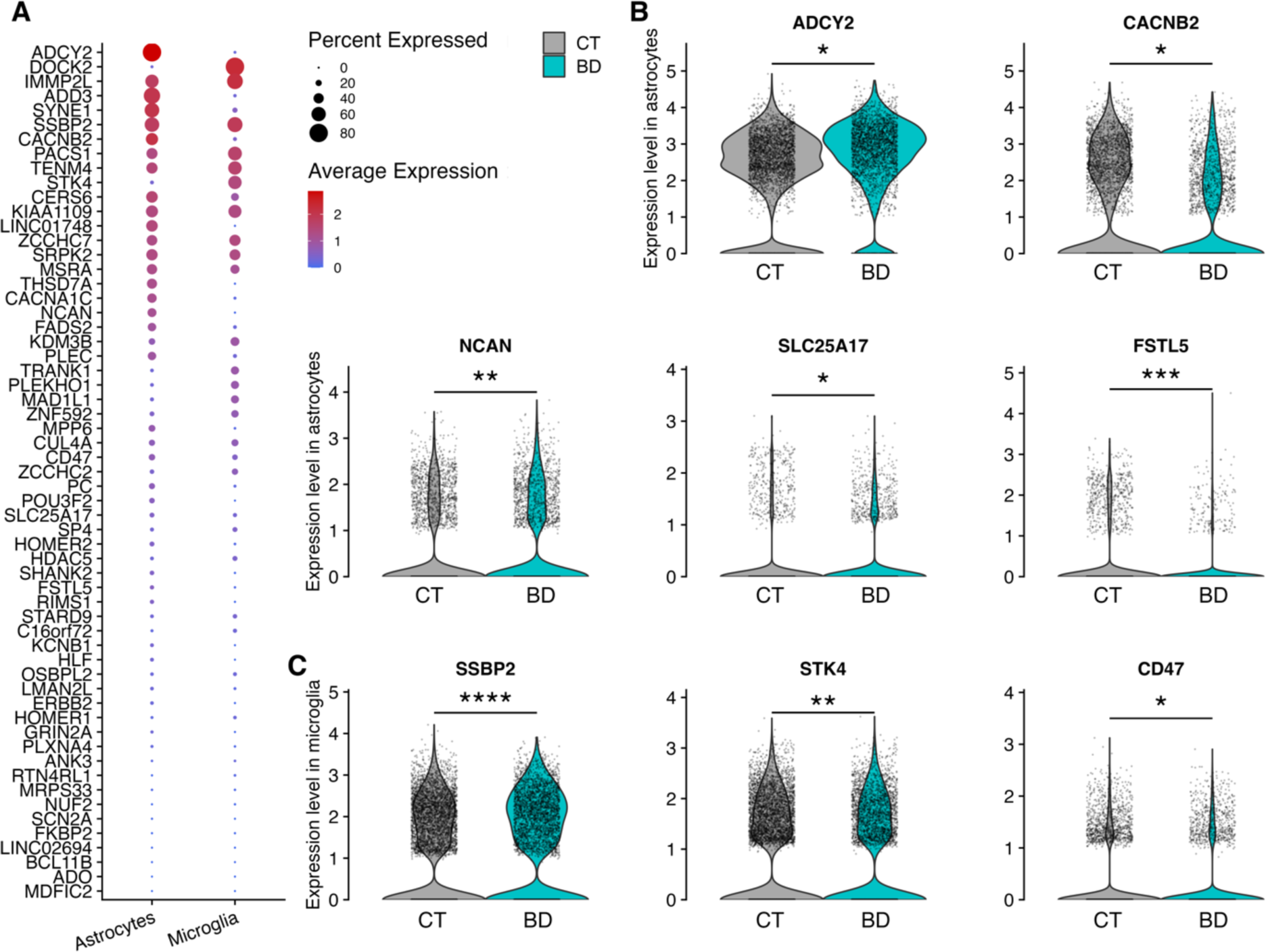
BD GWAS genes are expressed in microglia and astrocytes. (**A**) Expression of the BD GWAS genes identified by Stahl et al. and Mullins et al. in microglia and astrocytes. (**B**) Differential expression of the BD GWAS genes ADCY2 (padj = 0.0923, LogFC = 0.699), CACNB2 (padj = 0.0669, LogFC = −2.607444), NCAN (padj = 0.0151, LogFC = 0.785), SLC25A17 (padj = 0.0369, LogFC = 0.268) and FSTL5 (padj = 0.00704, LogFC = −0.577) in astrocytes of BD vs CT. (**C**) Differential expression of the GWAS genes SSBP2 (padj = 0.000185, LogFC = 0.755), STK4 (padj = 0.0231, LogFC = −0.150) and CD47 (padj = 0.0856, LogFC = −0.199) in microglia of BD vs CT. (correspondence of adjusted p value in graphs: *: p < 0.1, **: p < 0.05, *** p < 0.01, ****: p < 0.001)

### Microglia in BD are characterized by reduced proinflammatory signalling and increased apoptosis

Having demonstrated that the proteomic changes in BD involves microglial molecules and that GWAS genes are enriched in microglia, we then examined the transcriptomic evidence of microglial dysfunction in BD using snRNAseq. DGE analysis in microglial nuclei yielded 379 DEG that are significantly downregulated in BD and 448 DEG that are significantly upregulated (*p*adj < 0.1, |logFC| > 0.1) (Figure 4A). FEA showed an upregulation of Wnt β-catenin-independent signalling (*TNRC6C*, *PPP3CA*, *RYK*, *PSME4*, *AGO2*, *ITPR2*, *CTNNB1*, *PRKCA*) and insulin signalling (*MAP3K3*, *STXBP3*, *INSR*, *MINK1*, *INPPL1*, *IRS2*, *PRKCA*, *FOXO3*, *PIK3CA*, *TBC1D4*, *MAP3K4*, *MAP2K5*, *MAP3K5*) along with PI3K-Akt, MAPK and calcium signalling pathways (Figure 4B). Insulin signalling, along with the upregulated *CSF1R,* both potential activators of the PI3K-Akt pathway, may drive autophagy^59^, which also shows evidence of upregulation. Autophagy might be in response to oxidative stress, which characterises BD^60^, especially given the dysfunctional antioxidant responses and glutathione metabolism indicated by the downregulation of relevant genes (*GSR*, *MGST2*, *LAP3*, *JUN*, *PNPT1*, *SELENOS*, *CYBA*, *VRK2*, *ETV5;* Figure 4A). The deficient stress response may be further aggravated by lysosome and phagosome-related gene downregulation (*ATP6V0B*, *ASAH1*, *CTSZ*, *CD68*, *LIPA*, *IGF2R*, *CTSC*, *LGMN*, *HLA-DRB5*, *TUBA1B*, *HLA-DMA*, *HLA-B*, *CYBA*, *CALR*, *ACTB*, *ACTG1*, *HLA-E*, *HLA-DPA1;* Figure 4A) potentially leading to a cytoplasmic accumulation of autophagosomes. Microglial dysfunction is accompanied by a reduction in the expression of proinflammatory genes of the cytokine, antigen processing, interferon, and JAK-STAT pathways. This could have an impact on synaptic pruning by microglia impacting neuroplasticity and neuronal activity^61^ in BD. Finally, we found evidence for an increased apoptosis of microglial cells in BD. Indeed, several genes involved in antiapoptotic signalling were found downregulated (e.g. *TUBA1B*, *NAIP*, *YWHAG*, *YWHAQ*, *BCL2*, *BCL2L1, GLUD1* and *ACTG1*; Figure 4C)^62–68^. To confirm the alteration of these pathways on a larger scale, we performed a DGE analysis between BD and CT samples on a publicly available bulk RNA sequencing dataset, including over 200 samples from the Cg and the amygdala, published by Zandi et al.^15^. Using gene set enrichment analysis on the results of the DGE analysis, we showed the enrichment of many of the same pathways that were enriched in our snRNAseq analysis results (Figure S2A).

**Figure 4.**
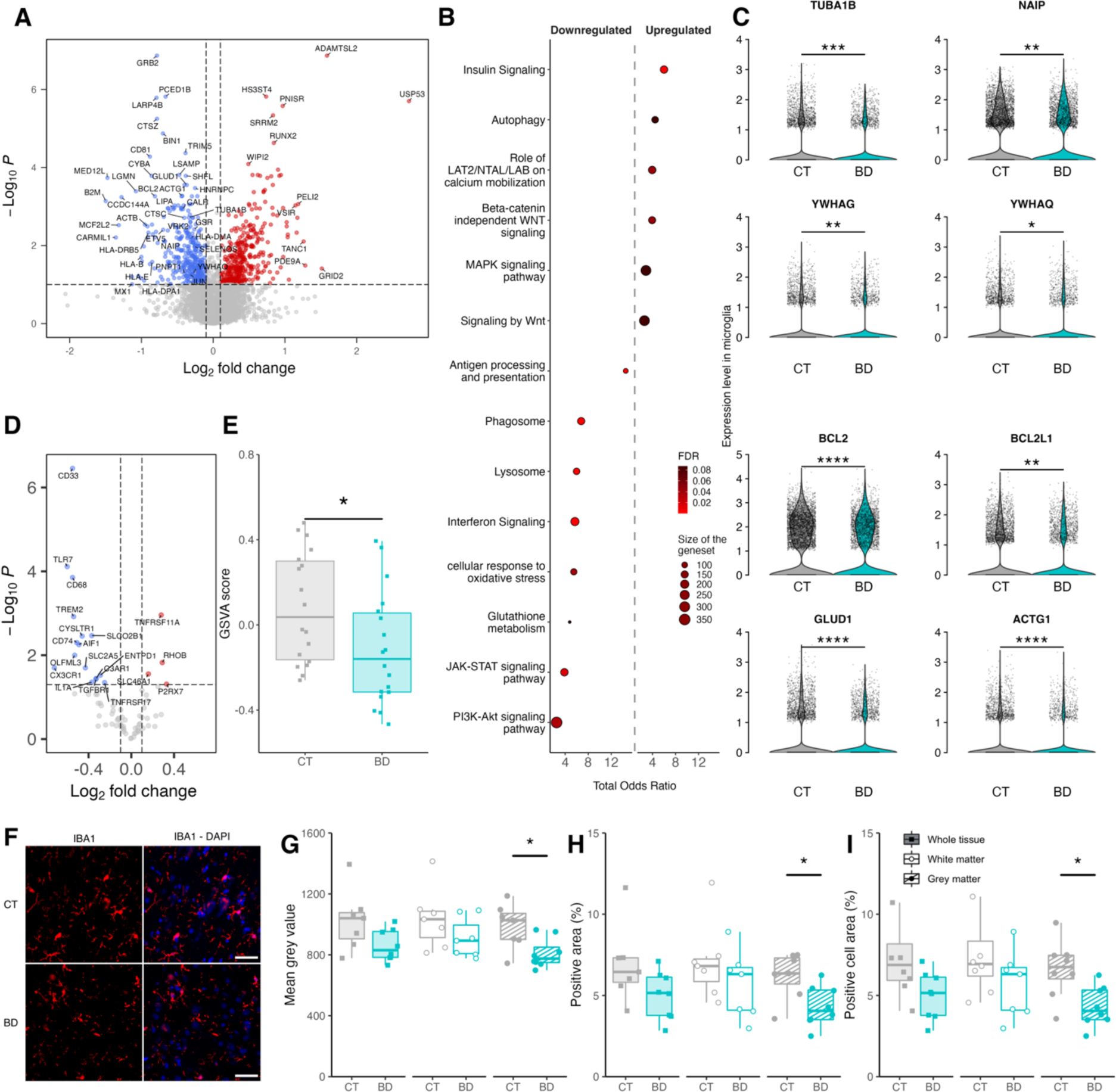
Microglial in BD shows lower proinflammatory gene expression along with increased apoptosis and reduced density in the Cg. (**A**) Volcano plot of the differentially expressed genes in microglia calculated with MAST. (**B**) Pathways enriched in microglial DEGs. (**C**) Expression of the antiapoptotic genes TUBA1B (p_adj_ = 0.00192, logFC = −0.3400), NAIP (p_adj_ = 0.01229, logFC = −0.4780), YWHAG (p_adj_ = 0.02118, logFC = −0.2563), YWHAQ (p_adj_ = 0.05241, logFC = −0.2763), BCL2 (p_adj_ = 0.0005483869, logFC = −0.81141), BCL2L1 (p_adj_ = 0.0156834605, logFC = −0.66311), GLUD1 (p_adj_ = 0.0002807306, logFC = −0.3679) and ACTG1 (p_adj_ = 0.0005346, logFC = −0.4303) in microglia in BD vs CT. (**D**) Volcano plot of the nCounter DEGs, only showing the microglia-specific genes highlighted by Butovsky et al. (**E**) GSVA score of the enrichment of the Butovsky microglial gene set in the the nCounter bulk transcriptomic dataset (t.test, p = 0.03057). (**F**) Representative image of the immunofluorescence staining of IBA1 (red) and nuclei (blue) on Cinglulate Cortex (Cg) of the CT (up) and BD (down). Scale bar = 50 µm. (**G**-**I**) Quantification of the mean intensity quantification of the IBA1 staining (**G**), IBA1 positive area (% of total tissue, **H**) and the number of detected IBA1 positive cells over total tissue area (cell/µm2, **I**) in the Cg of BD as compared with control in the whole tissue (WT, filled boxes, ▪), white matter (WM, blank boxes, ○) or grey matter (GM, dashed boxes, ●) (significance as the result of bilateral t-test, *: p < 0.05).

To further validate the snRNAseq results, we performed a bulk transcriptomic comparison between our BD and CT samples, using the nCounter Glial Expression Panel (Nanostring) supplemented with 55 custom genes (*Table S3*). This analysis was performed separately in tha grey (GM) and white matter (WM). When we jointly analysed both regions (whole tissue, WT), 52 genes were significantly downregulated and 40 upregulated (p < 0.05). When GM and WM were analysed separately, 36 significantly downregulated and 28 significantly upregulated genes were found in the GM (p < 0.05), whereas 58 significantly downregulated and 14 significantly upregulated genes were found in the WM (p < 0.05, Figure S2B-E). We tested the presence of a set of human microglia-specific genes highlighted by Butovsky et al.^31^ among the DEGs in the WT and separately in GM and WM. The majority of these genes were downregulated in BD (16 significantly downregulated for 4 significantly upregulated genes), with prominent examples of *TREM2*, *CD68*, *CD74*, *CD33*, *AIF1*, *TLR7* and *CX3CR1* (Figure 4D and Figure S2B-E) in all comparisons. We then performed a gene set variation analysis (GSVA)^69^ on the nCounter dataset, to compare the overall expression of microglia-specific genes^31^, which also showed a significant downregulation of this gene set in BD (Figure 4E). These results confirm a reduction of genes related to microglial proinflammatory and phagocytosis pathways. To test the reproducibility of this result on a larger scale, we performed the same analysis on the dataset published by Zandi et al. and found that, in addition to the validation of an overall hypoactivation of microglia in Cg in BD (Figure S2F), this result extends to the amygdala (Figure S2G).

With evidence of increased apoptosis in microglia in BD and a global downregulation of microglia specific genes in these two bulk transcriptomic datasets, we hypothesized that microglial density may be reduced with the disease. We thus performed IBA1 immunofluorescence (IF) assay (Figure 4F), which confirmed a significant decrease of IBA1 intensity (Figure 4G) and density (Figure 4H) in the GM (*p*_mean grey value_ = 0.02195, *p*_positive area_ = 0.01962). This decrease of intensity and density of IBA1 is accompanied by an *increase* of IBA1 positive cell number (Figure 4I) (*p*_positive cell area_ = 0.01054).

Given the age of the subjects in our study and the occurrence of Alzheimer’s disease (AD)-related pathology even in non-demented elderly subjects, we also asked if such pathological changes could bias the results of our study (e.g. if the higher proinflammatory and phagocytosis-related pathway enrichment in CT subjects was due to a possible higher level of AD-type pathology). We first assessed the presence of amyloid β deposits as well as hyperphosphorylated tau protein in both BD and CT subjects (Figure S3) that showed no significant difference. We next asked if the differential expression of microglial genes could have been induced by psychotropic medications that some of the BD subjects of our cohort were taking. We thus tested the overlap of DEGs from our microglial nuclei with known gene sets influenced by the treatment with lithium^23^, olanzapine^24^ and valproate^25^, without finding any significant overlap. This result confirms that microglial transcriptomic alterations in our dataset have not been the result of medication, but most likely of the disease itself. Overall, our results point towards a dysfunctional microglia phenotype in BD, showed by a decreased density, reduced phagocytic capacity and proinflammatory activation, associated to a deficient response to oxidative stress and apoptosis.

### Astrocytes in BD show transcriptomic alterations suggestive of metabolic and immune dysfunction and impaired extracellular matrix composition

After having established the impairment of microglia in BD and because the proteomic analysis of our samples also pointed towards astrocyte-specific alterations, we next looked for astrocyte-specific transcriptomic alterations in our snRNAseq dataset and performed a DGE analysis in these cells. It resulted in 288 genes significantly downregulated in BD and 501 significantly upregulated (*p*adj < 0.1) (Figure 5A). We found evidence for dysfunctional lipid metabolism in BD (Figure 5B). Fatty acid, triacylglycerol, and ketone bodies metabolism-related genes (*SLC27A1*, *ACADVL*, *SMARCD3*, *SCD5*, *ACOT11*, *NPAS2*, *ACSF2*, *AGPAT3*, *MED13*, *RXRA*, *SIN3B*, *PPARA*, *ACAD10*, *CLOCK*), including genes implicated in lipid transport and fatty acid oxidation in astrocytes, were upregulated. On the contrary, genes implicated in phosphatidylcholine and phosphatidylethanolamine metabolism and trafficking, important for mitochondrial and synaptic function^70–72^ were downregulated (*SLC44A3*, *CHKA*, *ABHD3*, *CHPT1*, *LPIN2*, *ETNK1*, *PLSCR1*, *PLSCR4*). Metabolic alterations are also supported by a decreased in insulin signalling pathway genes^73^ (*SPRED2*, *SPRED1*, *RAP1A*, *PSMA1*, *PSME4*, *PDE3B*, *IQGAP1*, *PPP2R5C*, *DUSP16*, *JAK1*), despite the significant upregulation of the insulin receptor in BD (*INSR*).

**Figure 5.**
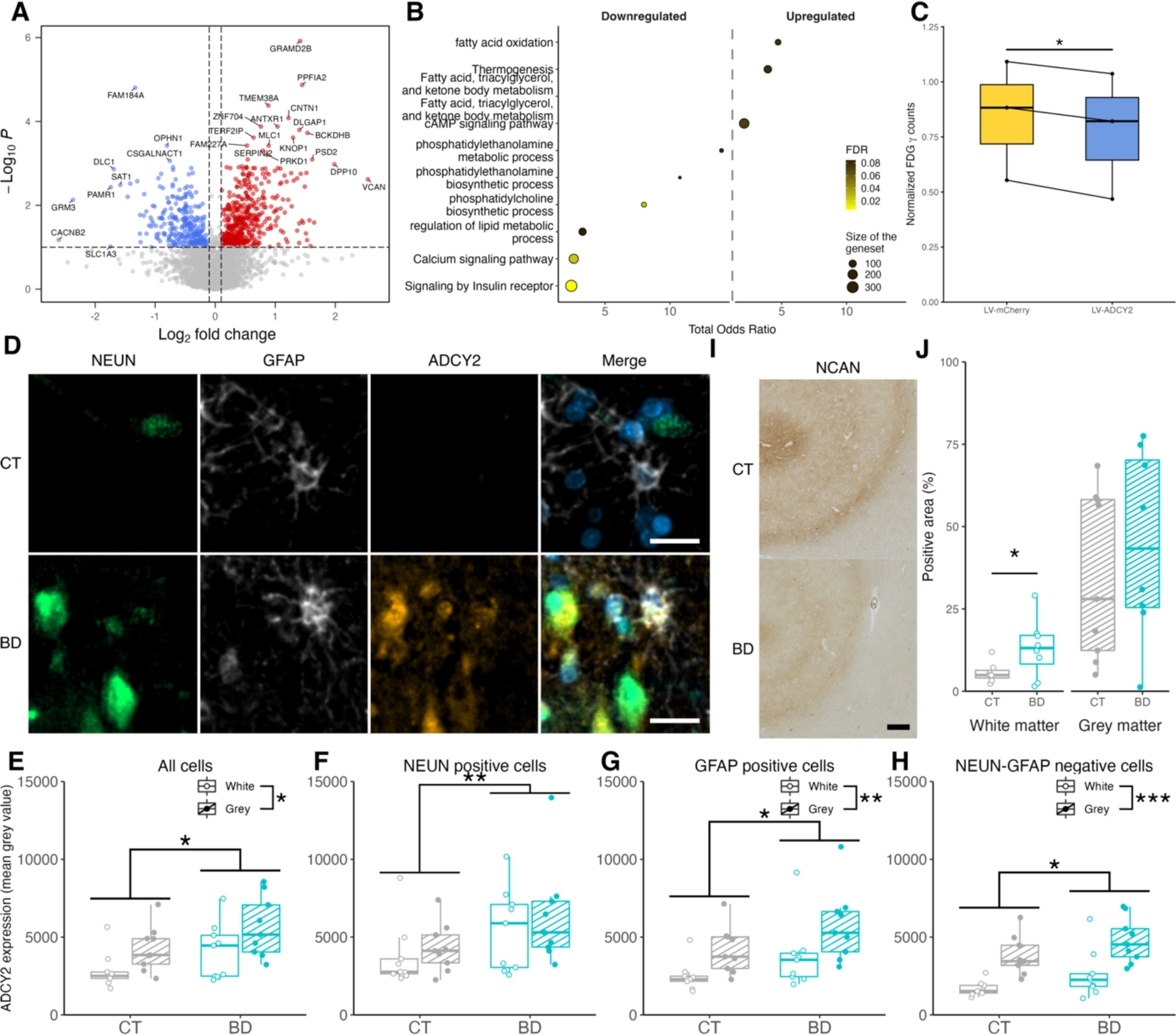
Overexpression of GWAS genes ADCY2 and NCAN in astrocytes is associated with altered metabolism, reduced proinflammatory gene expression and altered ECM composition. (A) Volcano plot of the astrocytic DEGs. (B) Pathways significantly modulated by the differentially expressed genes (DEGs) in astrocytes. (C) Result of the normalized quantification of [^18^F]FDG γ counting from cells infected with either LV-mCherry or LV-ADCY2 viruses. Paired t-test, p = 0.0188. (D) Representing images of the NEUN (green), GFAP (grey) and ADCY2 (orange) immunofluorescence staining in the Cg of BD and CT subjects. Scale bar = 20 µm. (E) Results of the quantification of the mean grey value of ADCY2 in various cell types. Results of a three-way anova with effect on diagnosis (p = 0.02) and region (p = 0.0107), but no interaction effect between those 2 parameters (p = 0.9059). (F) Results of the quantification of the mean grey value of ADCY2 in NEUN^+^ cells. Results of a three-way anova with effect only on diagnosis (p = 0.0282), but not on region (p = 0.3973). (G) Results of the quantification of the mean grey value of ADCY2 in GFAP^+^ cells. Results of a three-way anova with effect on diagnosis (p = 0.02503) and region (p = 0.00514), but no interaction effect between the 2 parameters (p = 0.7985). (H) Results of the quantification of the mean grey value of ADCY2 in GFAP^-^/NEUN^-^ cells. Results of a three-way anova with effect on diagnosis (p = 0.0278) and region (p < 0.005), but no interaction effect between the 2 parameters (p = 0.9289). (I) Representative image of NCAN immunohistochemistry staining in the Cg of CT (up) and BD (down) subjects. Scale bar = 250 µm. (J) Results of the quantification of NCAN-positive pixels over total pixel (in %) in the Cg of BD as compared with control (significance as the result of unilateral t-test, *: p < 0.05).

Calcium and cAMP signalling are important second messengers of signals involved in glucose and glycogen homeostasis and of lactate production and both these second messengers are modulated by adrenergic signalling. Our results show that the gene coding for the adrenergic α1 receptor (*ADRA1A*) was upregulated in BD, along with the downstream thermogenesis pathway genes (*SMARCD3, ADCY2, ARID1A, ADCY8, PRKG1, MAP3K5, SMARCA4, FGFR1*)^74^. However, genes implicated in the mobilisation and the increase of calcium and calcium signalling were downregulated (*PPP3CA, GNA14, PPP3CC, STIM2, ITPR2, PLCG1, CALM2, SLC8A1*), including the GWAS gene *CACNB2*, which also codes for a subunit of a voltage-gated calcium channel^39^. In parallel, we found an upregulation of cAMP producing enzymes *ADCY2* and *ADCY8,* with a downregulation of *UGP2,* which contributes to glycogen synthesis. These results suggest an imbalance between astrocytic cAMP- and Ca-dependent signalling pathways, which along with the downregulation of glycogen synthesis may contribute to deficient glucose and lactate homeostasis in the BD brain^75,76^. Several of these pathways were also found altered in the DGE analysis of the Zandi et al. dataset (Figure S2A) We then asked if upregulation of ADCY2^77^, which is also related to the genetic risk for BD may have a functional impact on astrocyte glucose metabolism. We overexpressed *ADCY2* in cultured primary mouse astrocytes and found a reduced [^18^F]-Fluorodeoxyglucose uptake by LV-ADCY2 infected astrocytes, as compared with mCherry-coding control virus-infected cells (Figure 5C). This result seems to support the downregulation of glycogen synthesis and degradation pathways found with FEA in snRNAseq (Table S4). The upregulation of *ADCY2* was validated in the BD brain at the protein level. Because *ADCY2* is also expressed in neurons, we included them in our analysis. Using triple immunostaining, we confirmed that GFAP-labelled astrocytes, NEUN-stained neurons, and NEUN-GFAP double negative cells, all significantly upregulated ADCY2 in BD (Figure 5D-H). This indicates that ADCY2 upregulation is widespread among the various brain cell types.

Apart from metabolic functions, a dysfunctional glutamate homeostasis by astrocytes, with downregulation of the metabotropic glutamate receptor 3 (*GRM3*) and the excitatory amino acid transporter (*SLC1A3*) were suggested by our results (*padj*_GRM3_ = 0.007411, log_2_FC_GRM3_ = −2.369, *padj*_SLC1A3_ = 0.09633, log_2_FC_SLC1A3_ = −1.744). Also related to glutamate homeostasis and the overall neuronal function, astrocytes additionally showed evidence of upregulation of ECM components (*VCAN*, *NCAN*, *B3GAT2*, *HS6ST1*, *GPC6*, *HS6ST3*), that form perineuronal nets in the brain, maintain glutamate homeostasis^78^ and the excitatory/inhibitory signalling balance^48–50^ and are a possible target of lithium, a mood stabilizer widely used for BD treatment^79^. Given the fact that *NCAN* expression is increased in astrocytes, and has been identified as a gene associated to the genetic risk for BD^7^ we also performed immunohistochemistry (IHC) within the Cg which confirm this upregulation of NCAN in the WM (one-tailed t-test, *p* = 0.02867, Figure 5I, J).

Consistent with the proteomic analysis at the whole tissue level, we also show a downregulation of Wnt signalling-associated genes (*PPP3C*, *TCF7L2*, *PPP3CC*, *CTNNB1*, *ROR1*, *TP53*, *LGR4*) and of genes that contribute to growth factor signalling and an upregulation of apoptosis genes. Finally, as in microglia, immune-related pathways, notably interferon signalling genes (*HERC5*, *RANBP2*, *DDX58*, *STAT2*, *MX1*, *EIF2AK2*, *PLCG1*, *XAF1*, *B2M*, *TRIM22*, HLA*-E*, *JAK1*) were significantly downregulated in BD (Table S4).

Given the dysregulation of apoptosis-related genes in astrocytes^80^, we performed a GFAP fluorescence staining on our tissue to assess astrocytic density and reactivity (Figure S4A), which showed no difference between BD and CT (Figure S4C-D). These results corroborate with our nCounter bulk RNA sequencing results, which showed no difference between CT and BD either when enriching for an astrocytic-specific gene set^32^ (Figure S4D-I). However, when we assessed the enrichment of this astrocyte-specific gene set on the bulk RNAseq dataset from Zandi et al.^15^ we did find a significantly reduced GSVA score showing a decrease of astrocyte-specific genes in Cg of BD (*p* = 0.011), but not in amygdala (*p* = 0.074). We also tested if the transcriptomic alterations in astrocytes could be induced by medication. As with microglia, no significant overlap was found between our DGE results in astrocytes and the gene sets associated to the BD treatments lithium^23^, olanzapine^24^ and valproate^25^.

Altogether, these results show an impact of the disease on important processes via which astrocytes support neuronal function along with a decreased immune signalling, with the GWAS genes *NCAN* and *ADCY2* as potential upstream regulators of these processes.

### Spatial transcriptomics distinguish astrocyte molecular alterations in the grey and white matter in BD

Given the morphological and functional differences of astrocytes between WM and GM^81^ and the fact that astrocyte alterations in BD have been reported to differ between the WM and the GM^82^, we next tested if the BD-associated molecular alterations in astrocytes were dependent on their localization^83,84^. We thus employed spatial transcriptomics and the Digital Spatial Profiling (DSP) to perform a gene expression analysis in GFAP-“enriched” regions of interest (ROI) in WM and GM (Figure 6A). After confirming that the most highly expressed genes in our ROI were indeed predominantly associated to astrocytes (using EWCE, Figure 6B), we then sought to determine the spatial origin of snRNAseq astrocytic subsets (Figure 6C). We performed subclustering of our astrocytes and identified three groups of cells. We then identified the marker genes that characterized each subcluster and calculated their GSVA scores in WM-GFAP-positive vs GM-GFAP-positive areas (Figure 6D). This resulted in an association of Astro 1 subcluster, predominantly enriched in neurotransmitter related pathways (associated to glutamate homeostasis pathways) with GM. On the contrary, Astro 2 and Astro 3 subclusters (associated to energy and lipid metabolism and immune signalling) are associated with WM (Figure 6E). These results suggested that astrocytic functions are spatially delimited between the GM and the WM and the BD-associated alterations differentially impact these two regions of the cortex.

**Figure 6.**
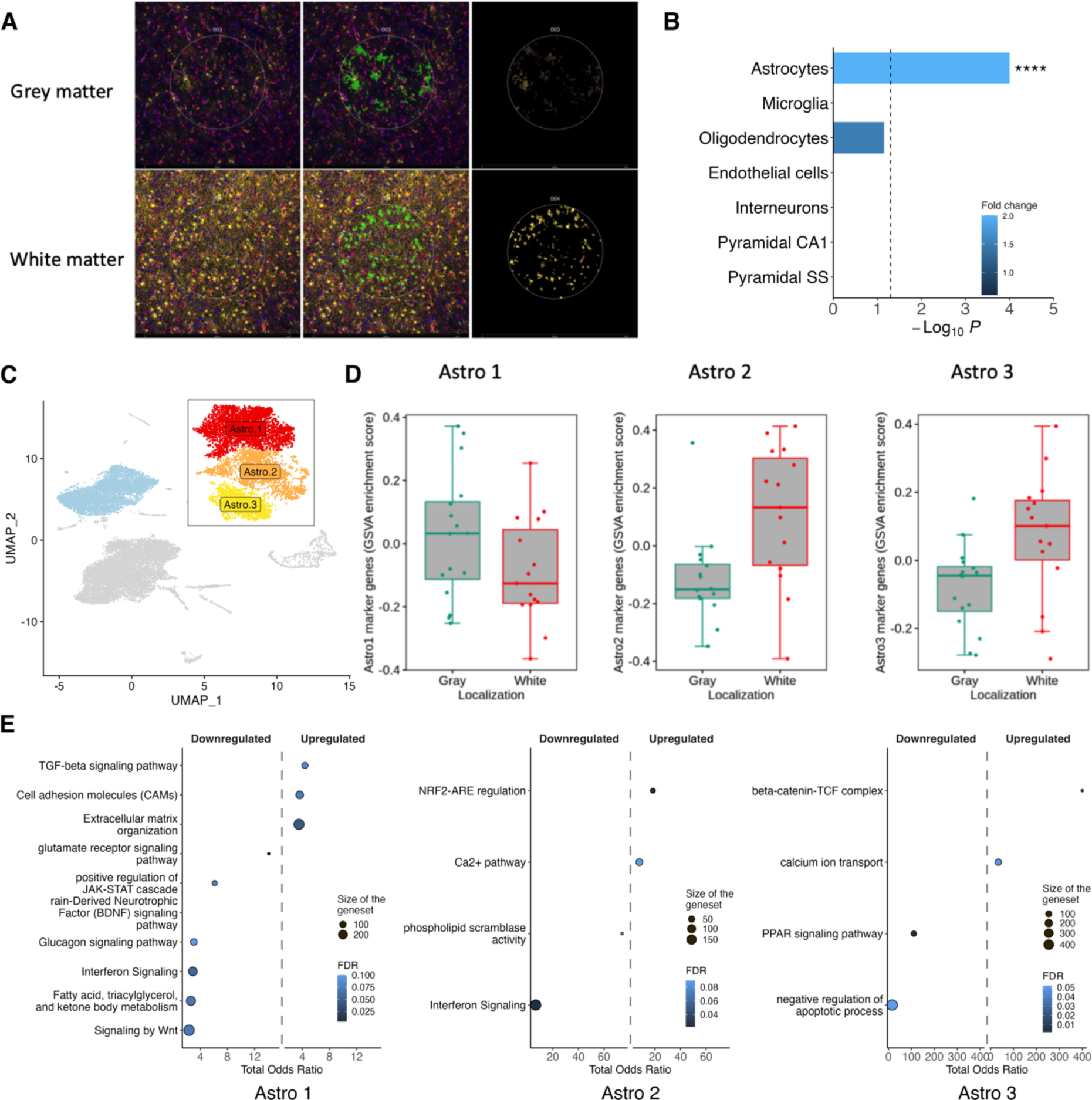
Spatial mapping of astrocyte dysfunctional pathways. (**A**) Representative image of the region of interest (ROI) localization in the GM and the WM (area of illumination (AOI) on GFAP staining for astrocytes selection). (**B**) EWCE results from the top 50 expressed genes of the GFAP positive AOI demonstrating that the sequenced ROI were enriched in astrocytic transcripts. (**C**) UMAP plot of the 3 astrocytic subclusters. (**D**) Results of the GSVA of the average expression profile of the 3 astrocyte subclusters with spatial transcriptomics in GM and WM. (**E**) Pathways significantly modulated by the DEGs in astrocytes subclusters.

### Cell-cell communication analysis prioritizes SPP1-CD44 signalling to astrocytes in BD

Having identified astrocyte- and microglia-specific changes in BD, we assessed for possible interactions between those two cell types. We thus performed a cell-cell communication analysis on snRNAseq data using the CellChat package in R^85^. This package highlights pairs of ligands and receptors that may be active in intercellular communication using snRNAseq data. CellChat results suggested that the *SPP1*-CD44 ligand-receptor pair showed stronger intercellular communication potential in BD than in CT. *SPP1*, a ligand expressed by microglia, astrocytes, perivascular macrophages, and oligodendrocytes communicates with astrocytes *via* the receptor *CD44*. We tested for evidence of co-regulated *SPP1*-*CD44* signalling in BD in our spatial transcriptomic results: indeed, GFAP-enriched ROI from the DSP experiment are *enriched* in astrocytic transcripts but also contain transcripts from immediately adjacent cells, other than astrocytes. Thanks to this very close spatial proximity, transcripts that show correlated expression across samples may thus indicate co-regulated gene sets. We calculated the correlation between CD44 and SPP1 expression separately in the CT and in the BD samples. We found a significant positive correlation only in BD. This result is an indication that this communication pair^86^ is more active in BD than in the CT brain (Figure 7A). Then, using co-staining, we investigated the presence of these 2 proteins in astrocytes at the tissue level (Figure 7B). We found that there were more GFAP^+^/SPP1^+^/CD44^+^ cells in the WM of BD than in CT (3-way ANOVA with Tukey post-hoc test, *p* = 0.0066, Figure 7C). To go further, we looked at the colocalization of those three proteins at the voxel level. We found that the colocalization of the three proteins is significantly increased in BD (3-way ANOVA, *p* = 0.0188, Figure 7D), further supporting an upregulation of SPP1-CD44 signalling in BD astrocytes. In light of our transcriptomic and IHC results, this upregulated communication appreas to be implicated in BD and could be contributing to the downregulated immune responses in astrocytes^87^ and dysfunctional glutamate homeostasis^88^.

**Figure 7.**
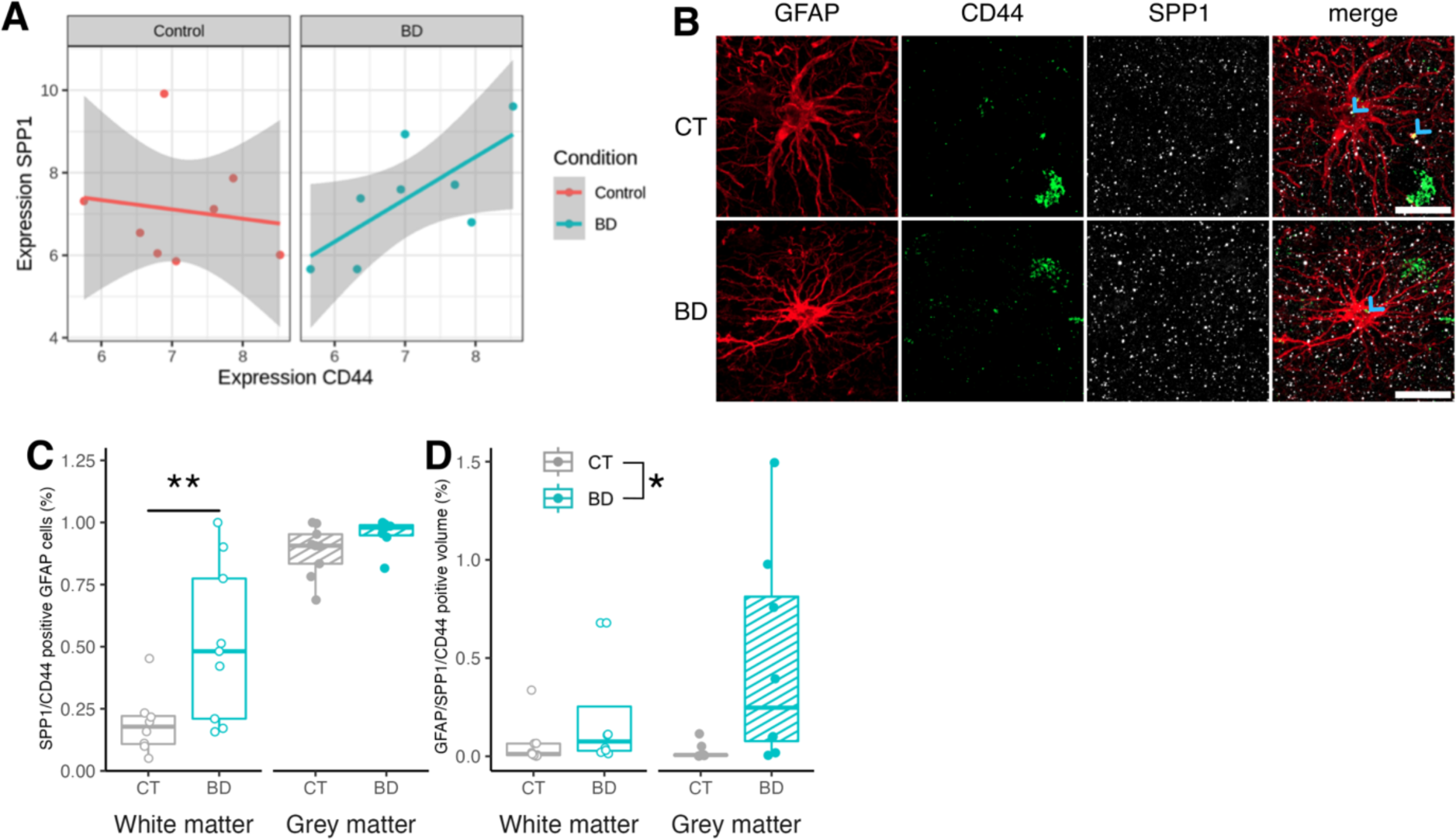
CD44/SPP1 communication is enhanced in BD. (**A**) Results of the expression of SPP1 and CD44 in the GFAP^+^ AOIs in spatial transcriptomics in BD and CT. (**B**) Representative image of the immunofluorescence staining of GFAP (red), CD44 (green) and SPP1 (white) in the Cq of BD and CT subjects. Blue arrows emphasise colocalization. Scale bar = 20 µm. (**C**) Result of the quantification of GFAP^+^/CD44^+^/SPP1^+^positive cells over total GFAP positive cells (%). Expression of Tukey post-hoc test after 3-way ANOVA: adjusted p = 0.0066. (**D**)Three-way ANOVA result of the quantification of GFAP^+^/CD44^+^/SPP1^+^ voxel as the percentage of total volume (%), diagnosis effect: p = 0.0188.

### Partial convergence between peripheral and central immune marker profiles indicates candidate serum biomarkers

Our overall results suggest a downregulated inflammatory response in microglia and astrocytes in BD. Nevertheless, increased peripheral inflammation in BD has consistently been reported. We used multiplex ELISA with a panel of 45 cytokines and immune system-related molecules (Table S5) on serum samples from an independent cohort of BD subjects and CT (n = 25 BD and n = 17 CT). We showed that several pro-inflammatory cytokines are increased in BD samples, such as IFNγ, IL-1b, IL-2 or TNFα. IL-10 and VEGF-D, which may have anti-inflammatory properties are also increased in BD (Figure S5A-P). Additionally, two anti-inflammatory cytokines were found decreased: LIF and HGF (Figure S5N-P). To test for associations between CNS cytokine expression and their peripheral concentration, we calculated the enrichment of genes coding for these cytokines in the Zandi et al.^15^ bulk RNAseq dataset using GSVA. The enrichment of genes coding for the proinflammatory cytokines that were upregulated in the serum of BD patients were unaltered in the brain. On the contrary, the genes coding for IL-10 and VEGF-D showed upregulated enrichment in the brain in BD, suggesting that unlike pro-inflammatory cytokines, the serum concentration of these two proteins may reflect their brain concentration (Figure S5Q-T). Taken together, our multiple ELISA results corroborate the widely described upregulation of proinflammatory cytokines in the serum of BD patients. However, this upregulation does not appear to originate in the brain.

## Discussion

In the present work, we have characterized molecular alterations in astrocytes and microglia in BD using a multi-omic approach on a cohort of *postmortem* human brain samples from the Cg. Bulk tissue proteomics provided evidence for downregulation of growth factor and proinflammatory signalling, citrate cycle and mitochondrial matrix proteins, accompanied by an upregulation of lipoprotein metabolism and apoptosis-related proteins. These significantly altered proteins were predominantly of astrocytic and microglial origin and this prompted us to perform a snRNAseq characterization of these two cell types in BD to obtain a cell-type specific mapping of the molecular alterations in the disease. We first showed that GWAS genes for BD were expressed in microglia and astrocytes and showed for the first time to our knowledge that several of them were altered with the disease (e.g. *ADCY2, NCAN, CD47, SSBP2)*. At the transcriptomic level, both microglia and astrocytes showed evidence for a downregulated proinflammatory and growth factor signalling and for dysfunctional metabolism. Using immunostaining we showed that microglial density is reduced in BD, associated with increased apoptosis and deficient response to oxidative stress. Among the potential mechanisms of astrocyte contribution to neuronal dysfunction, we found evidence for dysregulated immune and metabolic functions, dysfunctional glutamate homeostasis and an overexpression of ECM components that form perineuronal nets, alterations that are potentially associated to increased signalling through SPP1 to astrocytic CD44. Using spatial transcriptomics, we show that these astrocytic alterations differentially affect the cortical WM and GM. Finally, we indicate that despite the diverging proinflammatory molecular alterations in the CNS (downregulation) and the periphery (upregulation) in BD, two potentially anti-inflammatory molecules (IL-10 and VEGF-D) may serve as biomarkers of the disease as their serum dynamics parallel those of the brain.

We provided several lines of evidence that proinflammatory signalling is downregulated in both microglia and astrocytes in BD, a finding that has been suggested by previous brain transcriptomic studies^15,89^. This comes in contrast with the largely reproduced increased concentration of proinflammatory cytokines in the plasma^1,11,12^ of BD patients. This could suggest that the peripheral proinflammatory markers are increased due to brain-independent processes^90^. Alternatively, this finding could be due to a particular vulnerability of BD patient glial cells to the effect of ageing. Indeed, our sample included aged individuals, whereas studies measuring peripheral inflammation have been performed on cohorts of a younger age. We can thus hypothesize that glial cells, particularly microglia present a « dystrophic » phenotype^91–93^ with an altered morphology (explaining the smaller IBA1^+^ area per cell in our study), an increased apoptosis and a lower proinflammatory gene expression. This could be due to chronic activation^94^. It is still unclear if during an acute episode of the disease there is proinflammatory activation in the brain, which could contribute to this chronic activation with multiple disease episodes. Such dysfunctional microglia may lead to defective synaptic pruning^95^ and, together with impaired astrocytic glutamate homeostasis may contribute to synaptic dysfunction in BD^96^. This result prompts further research into carefully phenotyped cohorts to identify clinical trajectories associated with glial cell dysfunction in BD. We have recently shown that Positron Emission Tomography (PET) imaging measuring the 18 kDa-Transclocator Protein (TSPO) in the CNS is a marker of microglial density^97^. PET TSPO imaging is the appropriate tool to further explore the clinical correlates of microglial density alterations in BD. The only PET TSPO study to date in BD included patients from various mood states, thus the upregulation that was highlighted could have been driven by individuals experiencing a disease episode^13^. In addition, there may still be peripheral inflammatory markers that show variations consistent with those in the brain, such as IL-10 and VEGF-D so further research should validate these markers.

Metabolic dysregulation was prominent in our proteomic and transcriptomic datasets. Insulin signalling genes were found downregulated in both microglia and astrocytes, potentially underlining a mechanism through which central glucose metabolism is implicated in BD pathophysiology^98^. *ADCY2* upregulation *in vitro* reduced astrocytic glucose uptake. At the same time, *ADCY2* is a BD GWAS gene: it could thus be an upstream of these metabolic alterations in astrocytes. This is particularly relevant, given the evidence in favour of an imbalanced cAMP and calcium signalling and glycogen synthesis, with a potential impact on lactate metabolism and the overall cortical activity^99–101^. Clinical imaging studies have shown that glucose metabolism is altered in BD^102,103^ and our study, in accordance with literature, underlines the contribution of microglia and astrocytes in this process^104,105^. It is important to note that 5 of the patients in our cohort where diabetic, although they were equally distributed between the samples (n = 2 in BD, n= 3 in CT) so the downregulation of insulin pathway genes is therefore probably not the result of this comorbidity.

Our spatial transcriptomic experiment allowed us to obtain information on the phenotypic characteristics of cortical GM and WM astrocytes. This finding provides potentially mechanistic insight into the WM connectivity impairment^106^ and GM volume alterations in BD.

We recognize that our study has limitations. The principal one is the small group size and, though we have provided multiples lines of evidence using independent experimental approaches to validate our most important results (including the use of a particularly large bulk RNAseq dataset). In addition, several individuals of our BD cohort (3 out of 9) were on BD medication (lithium, clozapine and quetiapine) at the time of death. This could have an impact on some of our results, but the relative contribution of any of those individuals on the whole cohort should be small. We specifically tested for overlap between the genes that we were altered in our snRNAseq analysis and genes altered in response to these medications from previous studies. We did not find any significant overlap and this supports the argument that the results of our DGE analysis in microglia and astrocytes is likely not due to the effects of psychotropic medication. Furthermore, if the results of our DGE analysis were biased by medication, we would most likely expect an opposite effects on mechanisms such as brain proinflammatory activation^107^ than what our results suggest. Another limitation stems from the fact that We studied samples from one brain region and our results warrant further validation in other brain regions implicated in BD. Finally, knowing that BD and other severe psychiatric disorders, such as schizophrenia and major depressive disorder may share common pathophysiological mechanisms, our study is limited by the fact that it does not provide any information on the specificity of our findings in BD.

In summary, we have performed a multi-omic analysis of human brain samples in BD and identified potential mechanisms affecting microglial and astrocytic function. We show that brain inflammatory signalling is reduced in BD and highlight potential serum cytokines that could reflect their brain levels, while glucose and lipid metabolism alterations also characterise the disease. Our work prompts the assessment of the clinical impact of these alterations with metabolomic profiling of serum as well as through carefully designed clinical studies.

## Materials and Methods

### Human samples

Flash Frozen (FF) and Formalin Fixed Paraffin Embedded (FFPE) human *postmortem* cingulate cortex samples were obtained from the same 9 BD and 9 age- and sex-matched non-neurological and non-psychotic CT subjects. The brain samples and/or bio samples were obtained from The Netherlands Brain Bank (Netherlands Institute for Neuroscience, Amsterdam, www.brainbank.nl). All Material has been collected from donors for or from whom a written informed consent for a brain autopsy and the use of the material and clinical information for research purposes had been obtained by the NBB. Three of the BD subjects were treated with lithium of which, one was also taking quetiapine and lamotrigine, one bipolar subject was taking clozapine, and one was taking olanzapine with carbamazepine up to the last week prior to death. Serum from 25 BD and 17 age- and sex-matched non-demented CT subjects were collected and stored by the University Hospitals of Geneva. Detailed information can be found in Table 1 and *Table S6*. The exclusion criterion was the diagnosis of dementia and substance use disorder. All experimental procedures were performed with the agreement of the Cantonal Commission for Research Ethics (CCER) of the Canton of Geneva.

**Table 1.** Clinical details of postmortem human brain samples. Values are expressed as mean ± SD. All numeric values were compared with student t.test except for sex that was compared with Chi-squared test. Detailed information about subjects can be found in Table S6.

|  | CT | BD | *P value* |
| --- | --- | --- | --- |
| Age | 79.67 (± 11.8) | 74.44 (± 10.04) | 0.32696 |
| Sample age | 52.67 (± 9.61) | 55 (± 8.5) | 0.59965 |
| Sex (F/M) | 6F/3M | 6F/3M | 1 |
| Braak | 2.56 (± 1.42) | 1.83 (± 1.33) | 0.34163 |
| *Postmortem* Delay | 6.22 (± 1.48) | 7.73 (± 1.91) | 0.07944 |
| Weight | 1232.11 (± 107.26) g | 1175 (± 134.68) g | 0.33448 |

### Immunohistology

Brains were fixed in 10% formalin solution for 4 weeks before embedding in paraffin according to the NBB procedure. Twelve µm sections were cut on a microtome then dewaxed, rehydrated, treated for 10 minutes with 14 mmol/L Glycine, and used for immunohistology. For ADCY2 staining, sections were treated in a solution of Citrate-EDTA pH6.2 in a decloaking chamber for 25 min at 95°C, then treated with Akoya Biosciences TSA Blocking Buffer for 1 hour at room temperature before being treated according to Akoya Biosciences Tyramide Signal Amplification Cyanine 3 procedure. Primary antibodies were used as follows: ADCY2 (orb34050, Biorbyt – 1/20), GFAP (MA5-12023, Invitrogen – 1/250), NEUN (266 004, Synaptic Systems – 1/250). After PBS washing, secondary antibodies for GFAP and NEUN were used and amplification antibodies according to the kit were used for ADCY2. For CD44, SPP1 and GFAP staining, sections were placed in a decloaking chamber for 20 min at 95°C in a solution of citrate pH6.0. Primary antibodies were used as follows: GFAP (ab5541, Sigma-Aldrich – 1/100), CD44 (F10-44-2, abcam – 1/80) and SPP1 (ab63856, abcam – 1/100). Immunohistochemistry was performed for NCAN, amyloid β (Aβ) and hyperphosphorylated Tau (pTau) quantification. For NCAN, sections were treated in a solution of EDTA pH8 in a decloaking chamber for 25 minutes at 95°C, then incubated in 1xPBS + 0.5% Tritton X-100 + 3% Bovine Serum Albumin before adding the primary antibody (HPA036814, Sigma-Aldrich – 1/1000) for 48h at 4°C. For Aβ, sections were immerged in a 100% formic acid for 10 min followed by a brief wash in DI H2O and PBS and an incubation with anti-Ab (800701, Biolegend – 1/500) for 24h at 4°C. For pTau, sections were treated with a solution of 0.25% KMnO4 for 10 min followed by an incubation in a solution of 1% Alkaline phosphatase for 90 seconds. Slides were then incubated for 24h with anti-pTau primary antibody (MN1020, Thermo Fisher – 1/500). After PBS washing, the HRP secondary antibody was added for 1 hour at RT. The DAB (Sigma-Aldrich, 0.2 mg/ml) revelation of staining was performed in 1xPBS + H_2_O_2_ (100 µl/L). Sections were then treated in 0.01% Cresyl violet for 10 min for nucleus staining. All images were acquired using a Zeiss Axio Scan.Z1 (Carl Zeiss) with a 10x objective (10x / NA 0.45 Plan Apochromat), with a Hamamatsu Orca Flash 4 monochrome camera (0.65 µm/pixel) for fluorescence, or a Hitachi HV-F202FCL color camera (0.44 µm/pixel) for brightfield imaging. Confocal imaging was executed on an LSM800 Airyscan confocal microscope (Carl Zeiss) with a 40x objective (Plan-APO 40x/1.4 Oil DIC (UV) VIS-IR). Acquisition was done with Zen 2.3 (Carl Zeiss) software. Images were processed Qupath (version 4.3)^108^ or with the Fiji distribution of ImageJ^109^.

### Proteomic analysis

Tissues were flash frozen into a block by diving them into liquid nitrogen according to the NBB procedure. Punches were made into the blocks mostly in grey matter, then put in 0.1% RapiGest (Waters) in 100 mM TEAB (Sigma Aldrich) for ultrasonic tissue lysis and protein extraction. After centrifugation, supernatants were subjected to protein digestion using trypsin. Resulting peptides were analyzed by nanoLC-MSMS using an easynLC1200 (Thermo Fisher Scientific) liquid chromatography system coupled with an Orbitrap Fusion Lumos mass spectrometer (Thermo Fisher Scientific). Data extraction and directDIA analysis was performed with SpectroNaut v15 (Biognosys) using the Human reference proteome database (Uniprot). We performed a post-hoc exclusion of 4 of the samples based on their high heterogeneity observed by PCA plot (n = 2 CT and n = 2 BD), as compared with the 14 other samples. Further analysis was performed with R (4.2) with the limma^110^ package for differential protein abundance analysis, and weighted correlation network analysis (WGCNA) package^17,111^ for co-expression module analysis^112^.

### Nanostring nCounter

#### Sample preparation

Tissues were flash frozen into a block by using liquid nitrogen according to the NBB procedure. Punches were made into the grey and white matter. Tissues were immediately immerged in TRIzol® (Invitrogen) then chloroform was added to the solution, and the RNA extraction was then performed according to the RNeasy Micro kit. RNA quantity and quality was assessed with a 2100 Bioanalyzer Instrument (Agilent). Samples were then processed in the nCounter (Nanostring) according to manufacturer procedure using the Glial Profiling Panel supplemented with 55 more genes (*Table S3*).

#### Quality control and Differential Expression

Quality control and normalization were done on raw data with the NanoTube^113^ R package. Differential expression (DGE) was performed with the same package, referring to the Limma DGE method, integrated in the package and accounting for known cofounders, according to the following design: *∼ diagnosis + batch + age + sex*.

### Single nuclei RNA sequencing

#### Sample preparation

The nuclei isolation for 10X Genomics nuclei capture was performed according to the protocol explained by Krishnaswami et al.^27^, adapted by Smith et al.^28^. Briefly, punches on GM and WM were made into blocks of tissue. The punches were then placed into nuclease-free buffer and crushed in a Douncer to obtain a liquid solution of tissue extraction. After centrifugation, the nuclei were then stained with anti-SOX10 antibody (R&D Systems, AF2864, 1:250) + anti-Goat AF488 (Invitrogen, A11055, 1:1000) and anti-NEUN AF647 (abcam, ab190565, 1:500) antibodies. Nuclei were then stained with Hoechst 33342 (abcam, ab228551) just before the Fluorescence Activated Cell Sorting (FACS) procedure, done in a MoFlo Astrios (Beckman Coulter). Around 15’000 NEUN-AF647-/SOX10-AF488-‘double negative’ nuclei were selected and sorted for further processing. Nuclei were prepared according to 10X Genomics Chromium Next GEM Single Cell 3’ v3.1 protocol targeting 6000 nuclei and captured on a Chromium Controller (10X Genomics). The sequencing was performed on an HiSeq 4000 (Illumina), with a sequencing depth of 1 line per sample.

#### Processing of FASTQ files

Demultiplexing, alignment, barcode filtering and UMI counting of the FASTQ files were processed with the CellRanger pipeline v6.1.2 (10X Genomics) with the inclusion of introns, aligned on the GRCh38 genome reference. Only protein coding genes were retained for further analysis.

#### Quality control, nuclei integration, dimension reduction and clustering

Resulting feature-barcode matrices files were processed following Nextflow (version 22.04.0) pipeline nf-core/scflow (version 0.7.0dev)^114^. Briefly, this pipeline performed QC for each sample, with exclusion criteria of a minimum and maximum of feature per nuclei between 700 and 2500. Nuclei with more than 5% of mitochondrial transcripts were excluded. When performing the QC, we noticed that one sample (BD, n = 1) needed to be removed due to an abnormally low number of captured nuclei, all the subsequent analysis were thus done without this sample. Sample integration was done using the Liger method^29^ which is included in the pipeline. For this integration, the optimal k value was found to be 30 and the optimal lambda value was found to be 5. Integration threshold value was set to 0.0001 and the maximum number of iterations was set to 100. Integrated nuclei were then clustered with the UMAP method^30^. Automated cell type annotation was performed following the nf-core/scflow pipeline, according to the prediction given with the use of the expression weighted cell type enrichment (EWCE) package^26^. Data were then processed with the Seurat suite of packages^115^ in R.

#### Differential gene expression analysis

Differential gene expression (DGE) analysis was performed separately in astrocyte and microglia clusters using the MAST^34^ package, which fits a zero inflated, negative binomial mixed effects statistical model using the following model: diagnosis+sex+batch+1|individual, where individual was the random effects variable.

#### Cell communication analysis

Cell communication analysis was performed on astrocytes and microglia using the CellChat^85^ package in R. This package predicted CT-or BD-specific as well as incoming-outcoming-, shared-, or incoming & outcoming-specific pathways, predicting expression of pairs of ligands and receptors.

### Nanostring GeoMx® Digital Spatial Profiler (DSP) Spatial transcriptomics

#### Sample preparation

Brains were fixed in 10% formalin solution for 4 weeks before embedding in paraffin according to the NBB procedure. Slide preparation and immunostaining were done according to Nanostring procedures, briefly: five µm section were cut followed by a Heat-Induced Epitope Retrieval at 100°C during 20 min in BOND Epitope Retrieval 2 (ER2) solution (Leica) with 0.1 Proteinase K before Deparaffinizing. Morphology marker staining was performed as the following: SYTO13 for nucleus staining (Nanostring), IBA1 (Millipore, MABN92-AF647, 1:100), GFAP (Novus Biologicals, NBP233184DL594, 1:300). In Situ Hybridization was then performed overnight. Finally, slides were placed in the GeoMx DSP. Regions of interest (ROI) were drawn as a circle of 650 µm of diameter in both grey and white matter. In each of the ROI, areas of illumination (AOI) were set. In the GM 2 AOIs per ROI were set, one for GFAP^+^ areas and one for IBA1^+^ areas. In the WM, AOIs were only set around GFAP^+^ areas. AOI selection was done with default parameters, except for the N-dilate parameter that was set to 3. AOI selection was carried out in such a way as to select only either GFAP^+^/IBA1^-^ or IBA1^+^/GFAP^-^ areas. After the selection of AOIs, the chosen areas were UV-micro-dissected according to the machine parameters and the transcripts thus released were recovered and whole transcriptome-sequenced according to Nanostring procedures.

#### Quality control

The resulting DCC and PKC files were proceeded with the help of the NanoStringNCTools, GeomxTools and GeoMxWorkflows packages with R. The default parameters were set for quality control filtering of the segments except for the minimum of read stitched that was set to 70%, the minimum of reads aligned to 65%, the minimum of negative control counts to 1, the maximum counts observed in NTC well to 25000, the minimum of nuclei estimated to 8 and the minimum of segment area to 1000. Then, genes that were not expressed by at least 10% of the segments were subtracted. Finally, gene expression was normalized with the quartile 3 (Q3) method.

### Bulk RNA sequencing

Previously published and publicly available bulk RNA sequencing data published by Zandi et al. (2022)^15^ were obtained from PsychENCODE Consortium on the NIMH Repository via Synapse under the BipSeq study (<u>syn5844980</u>). This study was made on a total of 511 samples from 295 individuals (138 cases and 157 controls) across the two brain regions: amygdala (n = 121, 121 BD and 122 CT) and subgenual anterior cingulate cortex (aCG, n = 268, 126 BD and 142 CT). Data was processed using the rna-seq nextflow pipeline^116^. The DGE analysis was performed using the limma-voom method, from the limma R package^110^ contained in the edgeR pipeline^117^. Gene set variation analysis was performed using the GSVA package in R, as previously described^118^.

### Functional enrichment analysis

Functional enrichment analysis (FEA) was performed using the enrichR^18^ package in R. Several functional enrichment databases were consulted including Wikipathways_2021_Human, KEGG_2021_Human, GO_Cellular_Component_2023, GO_Molecular_fonction_2023, GO_Biological_Process_2023, Reactome_2022 or BioPlanet_2019, using the significantly differentially expressed genes (DEG). FEA was also performed using the fgsea^119^ package in R.

### [^18^F]-FDG uptake on primary astrocyte

#### ADCY2 vector

Human *ADCY2* cDNA sequence (NM_020546.3) was fused to the 14 amino acids V5 tag^120^ at the 5’ end and placed under the control of a *GFAP* promoter. A Green Fluorescent Protein (GFP) coding sequence was placed under ubiquitous hPGK promoter as a viral infection control. Same lentiviral construction was made with mCherry construction in place of *ADCY2* sequence as control. The DNA sequence was synthetized by GeneArt Gene Synthetize (ThermoFisher) transfected with JetPRIME (Polyplus) in pCLX-EF1-GFP vector plasmid with psPAX2 packaging plasmid and pCAG-VSVG envelope plasmid and cloned in 293T cells. The full sequence can be found in Table S7. Lentiviral vectors have been produced by the Vector Lab core facility of the University of Geneva.

#### Cell culture

Primary mouse astrocytes were produced and given by GliaPharm (Geneva, Switzerland). Astrocytes were cultured in Poly-L-Ornitine pre-coated flat bottom 12-well plate in a high glucose DMEM (Sigma, D7777) medium supplemented with 44 mM NaHCO_3_ (Sigma, S4019), 0.01% Antibiotic antimycotic 100x (Sigma, A5955) and 10% FCS at 37 °C with 5% CO_2_. Cells were incubated during seven days prior either LV-ADCY2 or control LV-mCherry infection at 1.8×10^4^ transducing unit per well (TU/well). Three days after infection, medium was replaced by fresh virus-free D7777 medium. Seven days after infection, D7777 medium was replaced by glucose-free DMEM (Sigma, D5030) medium with 44 mM NaHCO_3_ (Sigma, S4019) and 2mM D-Glucose (Sigma, G7021).

#### [^18^F]-FDG incubation

Two days after Glucose deprived D5030 medium incubation, cells were incubated with 0.037 MBq per well of [^18^F]-Fluorodeoxyglucose (FDG) during 25 min, then immediately placed on ice to stop the reaction, washed with 4°C sterile PBS and harvested with 10 mM NaOH (Merck), 0.1% Triton X-100 (Sigma, 93426) supplemented with proteases/phosphatase inhibitors (Thermo Scientific, A32959), before gamma counting using a gamma counter (Wizard 3, PerkinElmer). [^18^F]-FDG uptake was normalized by quantity of protein for each well.

### Multiplex cytokines ELISA

Multiple target ELISA was performed on frozen serum sample from BD and age- and sex-matched CT subjects, using antibody-coated magnetic beads to determine the abundance of inflammatory cytokines, chemokines and growth factors, for a total of 45 different targets (EPX450-12171-901, Invitrogen). The complete list of targets can be found in Table S5. Samples were diluted 1:100 in the buffer provided in the kit (1X Universal Assay Buffer) and procedure was performed according to user guide (revision A.0). Plate was read using a MAGPIX (Luminex) xMAP instrument. Data analysis was done with the free online software ProcartaPlex Analysis App (Thermo Fisher) using the 5PL algorithm, and statistical analysis was performed uising ProcartaPlex nonparametric M-statistics algorithm for group comparisons.

## Supplementary data

*Table S1 – Proteomic results of the WGCNA module coexpression analysis*.

*Table S2 – Astrocytic and microglial expression of the GWAS genes identified by Stahl et al.*^7^ *and Mullins et al.*^10^.

*Table S3 – Set of genes used for nCounter experiment composed of the Glial Profiling Panel and 55 supplemented genes*.

*Table S4 – snRNA seq results containing markers for all clusters (and astrocytes subclusters), DEG for astrocytes and microglia as well as FEA results for astrocytes, astrocyte subclusters and microglia*.

*Table S5 – Multiplex 45-target ELISA results of serum samples*.

*Table S6 – Metadata of all the cohorts of subjects samples used throughout the experiments. Table S7 – Sequence of the ADCY2 LV-ADCY2 Lentiviral construction*.

## Acknowledgments

This work was supported by the Swiss National Science Foundation (no. 320030-184713 and no. 32003B_156914). Author ST was supported by a Nanostring Young Investigator Award, which funded the nCounter and the spatial transcriptomics experiment. Bio-samples and/or data for this publication were obtained from NIMH Repository & Genomics Resource, a centralized national biorepository for genetic studies pf psychiatric disorders. Data were generated as part of the PsychENCODE Consortium, supported by: U01DA048279, U01MH103339, U01MH103340, U01MH103346, U01MH103365, U01MH103392, U01MH116438, U01MH116441, U01MH116442, U01MH116488, U01MH116489, U01MH116492, U01MH122590, U01MH122591, U01MH122592, U01MH122849, U01MH122678, U01MH122681, U01MH116487, U01MH122509, R01MH094714, R01MH105472, R01MH105898, R01MH109677, R01MH109715, R01MH110905, R01MH110920, R01MH110921, R01MH110926, R01MH110927, R01MH110928, R01MH111721, R01MH117291, R01MH117292, R01MH117293, R21MH102791, R21MH103877, R21MH105853, R21MH105881, R21MH109956, R56MH114899, R56MH114901, R56MH114911, R01MH125516, and P50MH106934 awarded to: Alexej Abyzov, Nadav Ahituv, Schahram Akbarian, Alexander Arguello, Lora Bingaman, Kristin Brennand, Andrew Chess, Gregory Cooper, Gregory Crawford, Stella Dracheva, Peggy Farnham, Mark Gerstein, Daniel Geschwind, Fernando Goes, Vahram Haroutunian, Thomas M. Hyde, Andrew Jaffe, Peng Jin, Manolis Kellis, Joel Kleinman, James A. Knowles, Arnold Kriegstein, Chunyu Liu, Keri Martinowich, Eran Mukamel, Richard Myers, Charles Nemeroff, Mette Peters, Dalila Pinto, Katherine Pollard, Kerry Ressler, Panos Roussos, Stephan Sanders, Nenad Sestan, Pamela Sklar, Nick Sokol, Matthew State, Jason Stein, Patrick Sullivan, Flora Vaccarino, Stephen Warren, Daniel Weinberger, Sherman Weissman, Zhiping Weng, Kevin White, A. Jeremy Willsey, Hyejung Won, and Peter Zandi.

## Data availability

All data generated in this manuscript will be made available on the Yareta repository upon publication https://doi.org/10.26037/yareta:gust4l5k5je5vkwcmqbx2r6gzu.

## Code availability

Analysis scripts used in this manuscript will be made available on GitLab upon publication.

## Author contributions

QA, BBT, KC, ST, and PM designed the experiment. QA and BBT performed the experiment. CP provided serum samples; SL provided material for the *in vitro* primary astrocytes study; NF, AS, DRO, and YYL contributed to experimental design data analysis; VS provided material for the multiplex ELISA assay. QA, ST, BBT, AMB, LMM, KC, PM analysed and interpreted the data. QA and ST drafted the manuscript, and all the co-authors revised it.

---

**Figure S1.**
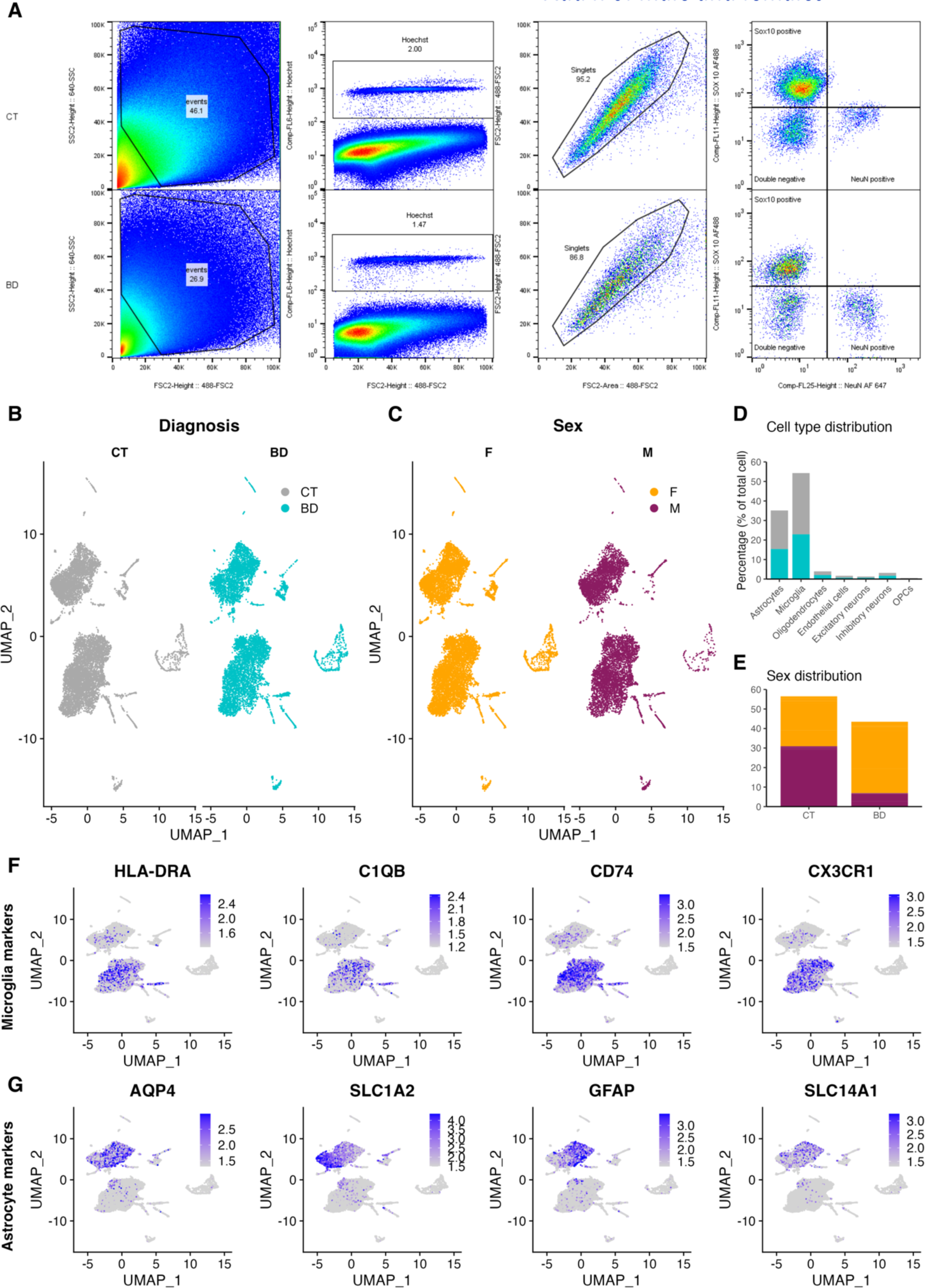
Detailed characterization of the snRNAseq results. (**A**) Representation of the negative nuclei selection by Fluorescence Activated Nucleus Sorting (FACS). Nuclei were first sorted by their shape and height, then by Hoechst-positive staining. Singlets were then selected, and the population of double Sox10- and NeuN-negative nuclei (= double negative, average = 16.98%, no difference between CT and BD, p = 0.496) was selected for further capture and single nucleus RNA sequencing analysis. (**B-C**) UMAP distribution of the nuclei respecting to the diagnosis (**B**) and the sex (F = Female, M = Male, **C**). (**D**) Distribution of every cell captured in snRNAseq according to their identified cell-type and subject diagnosis. (**E**) Distribution of every cell captured in snRNAseq by sex and diagnosis. (**F-G**) Expression plot of several microglia-specific (**F**) or astrocytes-specific markers (**G**).

**Figure S2.**
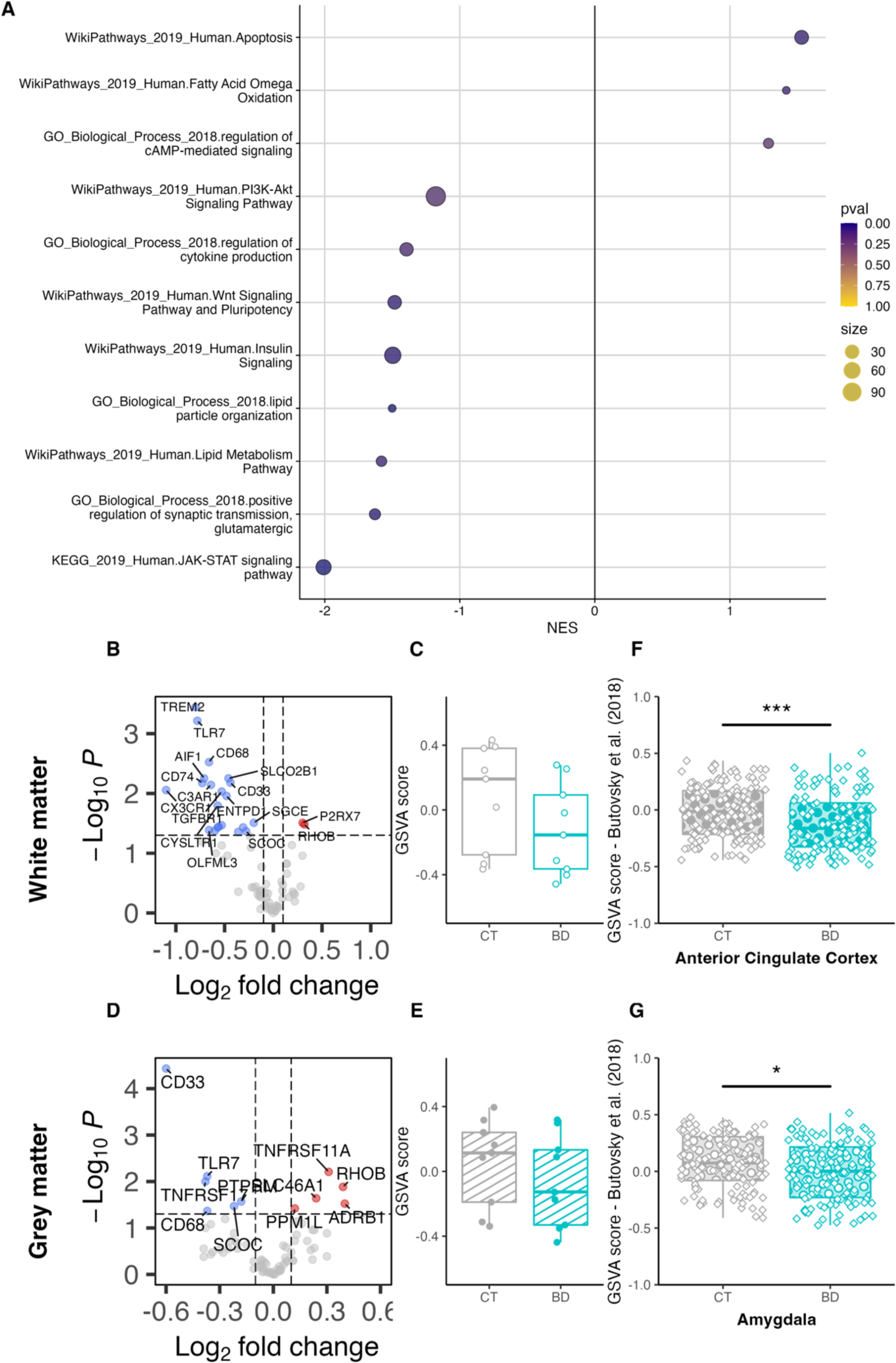
GSVA scores of microglial gen set in the nCounter results and bulk RNA seq from Zandi et al.^15^. (**A**) Representative pathways that were significantly enriched in the DEGs in the Zandi et al. bulk RNAseq dataset. (**B**) Volcano plot of the nCounter DEG in the white matter, only showing the microglial genes highlighted by Butovsky et al. (**C**) GSVA score of the enrichment of the Butovsky microglial gene set and the DGE from the nCounter experiment in the WM (t.test, p = 0.1929). (**D**) Volcano plot of the nCounter differentially expressed genes in the grey matter, only showing the microglial genes highlighted by Butovsky et al. (**E**) GSVA score of the enrichment of the Butovsky microglial gene set and the DGE from the nCounter experiment in the grey matter (t.test, p = 0.3833). (**F**) GSVA score of the DEG from Zandi et al. in the anterior cingulate cortex of BD and CT subjects with the curated microglial gene set from Butovsky et al. (t.test, p < 0.001). (**F**) GSVA score of the DEG from Zandi et al. in the amygdala of BD and CT subjects with the curated microglial gene set from Butovsky et al. (t.test, p = 0.01976).

**Figure S3.**
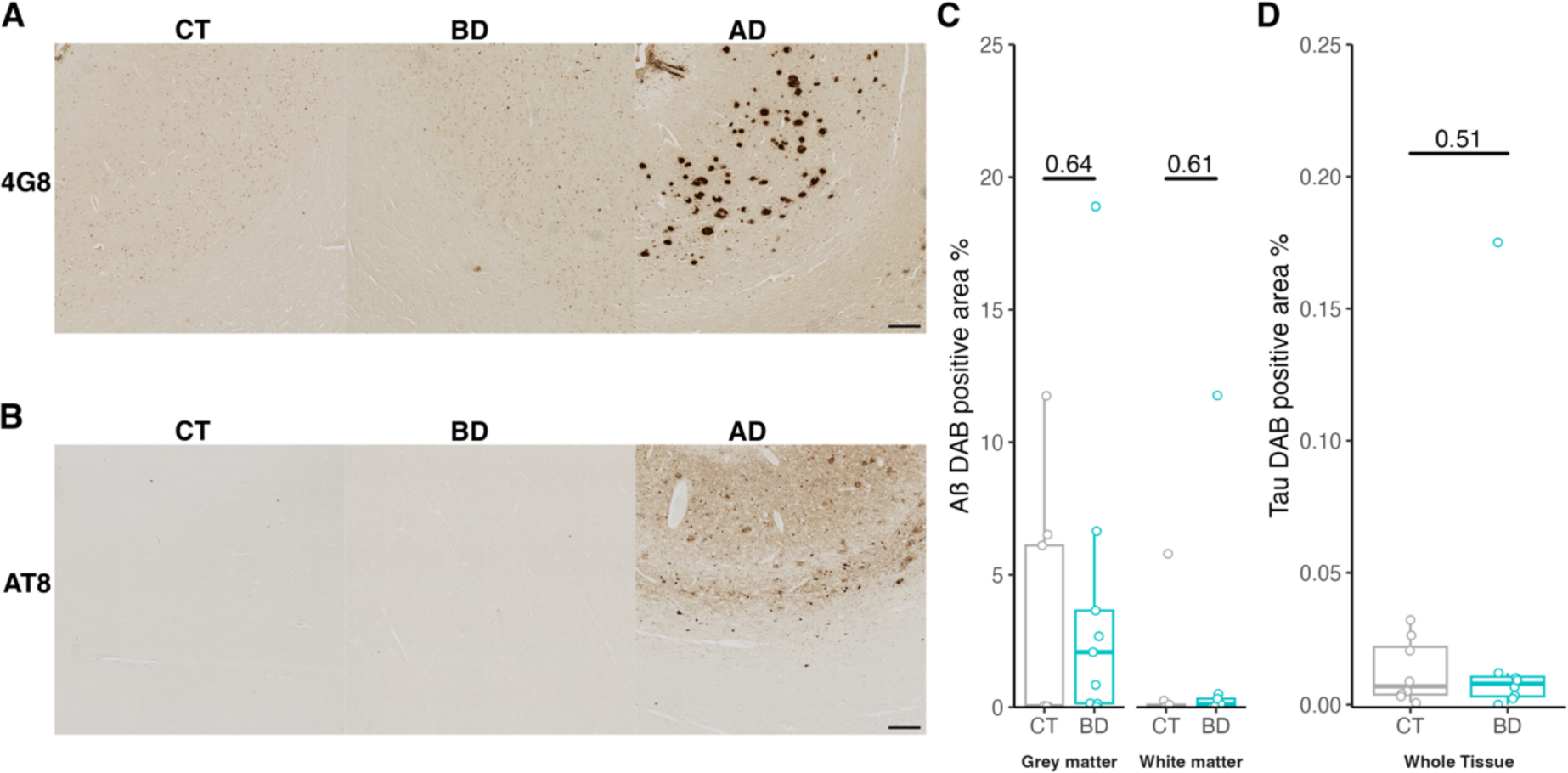
Evaluation of amyloid and Tau pathology in our cohort. (**A**) Representative image of the amyloid β specific 4G8 IHC staining of CT, BD and an AD case as positive control. (**B**) Representative image of the phosphorylated Tau specific AT8 IHC staining of CT, BD and an AD case as positive control. (**C**) Results of the quantification Amyloïd β-positive pixels over total pixel (in %) in the GM and WM of Cg in BD subjects compared to CT (expression of the result of student t-test). (**D**) Results of the quantification hyperphosphorylated Tau-positive pixels over total pixel (in %) in the Cg of BD subjects compared to CT (expression of the result of student t-test). AD case is shown as positive control example.

**Figure S4.**
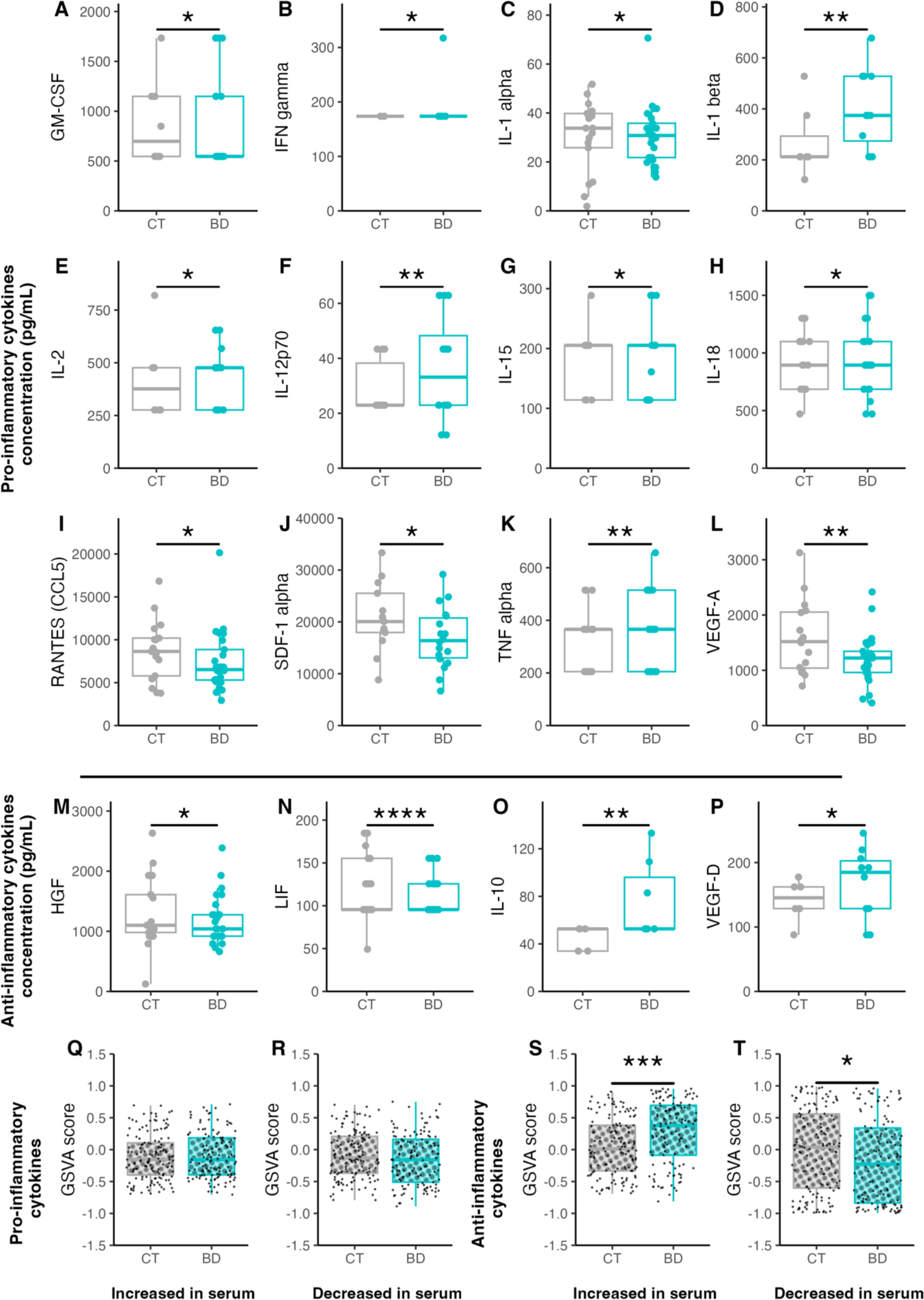
IF results of GFAP staining and GSVA scores of microglial gene set in the nCounter results and bulk RNA seq from Zandi et al. (**A**) Representing image of the GFAP staining. (**B**) Mean intensity quantification of the GFAP staining in the Cg of BD as compared with CT in the whole tissue (WT, filled boxes, ▪), white matter (WM, blank boxes, ○) or grey matter (GM, dashed boxes, ●) (significance as the result of bilateral t-test, *: p < 0.05). (**C**) GFAP positive area (% of total tissue) quantification in the Cg of BD as compared with CT in the whole tissue (WT, filled boxes, ▪), white matter (WM, blank boxes, ○) or grey matter (GM, dashed boxes, ●) (significance as the result of bilateral t-test, *: p < 0.05). (**D**) Quantification of the area of the detected GFAP^+^ cells over total tissue area (cell/µm^2^) as compared with CT in the whole tissue (WT, filled boxes, ▪), white matter (WM, blank boxes, ○) or grey matter (GM, dashed boxes, ●) (significance as the result of bilateral t-test, *: p < 0.05). (**E**) Volcano plot of the nCounter differentially expressed genes, only showing the astrocytic genes highlighted by Zhang et al. in the whole tissue. (**F**) GSVA score of the enrichment of the Zhang et al astrocyte-specific gene set and the DGE from the nCounter experiment in the whole tissue (t.test, p = 0.8446). (**G**) Volcano plot of the nCounter differentially expressed genes in the white matter, only showing the astrocytic genes highlighted by Zhang et al. (**H**) GSVA score of the enrichment of the Zhang astrocytic gene set and the DGE from the nCounter experiment in the white matter (t.test, p = 0.4921). (**I**) Volcano plot of the nCounter differentially expressed genes in the grey matter, only showing the astrocytic genes highlighted by Zhang et al. (**J**) GSVA score of the enrichment of the Zhang astrocytic gene set and the DGE from the nCounter experiment in the grey matter (t.test, p = 0.5863). (**K**) GSVA score of the DEG from Zandi et al. in the anterior cingulate cortex of BD and CT subjects with the curated astrocytic gene set from Zhang et al. (t.test, p = 0.07423). (**L**) GSVA score of the DEG from Zandi et al. in the amygdala of BD and CT subjects with the curated microglia-specific gene set from Butovsky et al. (t.test, p = 0.01139). (**M**) Volcano plot of the proteomic differentially abundant proteins highlighting the ACAN protein.

**Figure S5.**
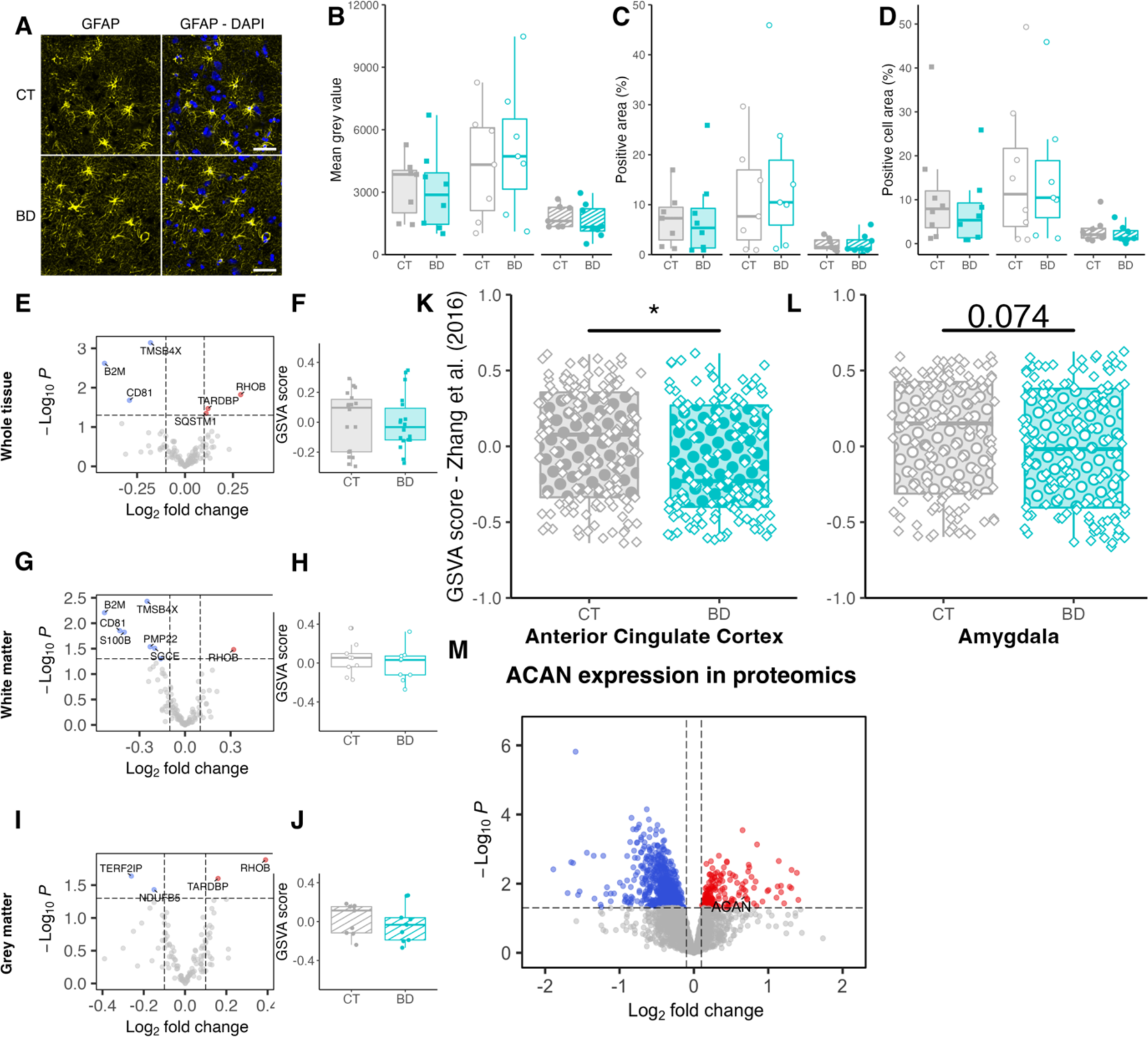
Multiplex ELISA results on the 45-plex human cytokines panel identifies serum cytokine alterations that parallel those of the CNS. (A-L) Significantly different (p < 0.05) results of the comparisons of serum concentration (in pg/mL) of pro-inflammatory cytokines in CT (n = 17) and BD subjects (n = 25). (M-P) Significantly different (p < 0.05) results of the comparisons of serum concentration (in pg/mL) of anti-inflammatory cytokines in CT (n = 17) and BD subjects (n = 25). Results are the expression of the nonparametric M-statistics algorithm for group comparisons. (Q) GSVA score of the DEG from Zandi et al. in the anterior cingulate cortex of BD and CT subjects with the significantly increased pro-inflammatory cytokines in the serum of BD subjects (n = 25) as compared with CT (n = 17), (t.test, p = 0.6899). (R) GSVA score of the DEG from Zandi et al. in the anterior cingulate cortex of BD and CT subjects with the significantly decreased pro-inflammatory cytokines in the serum of BD subjects as compared with CT, (t.test, p = 0.188). (S) GSVA score of the DEG from Zandi et al. in the anterior cingulate cortex of BD and CT subjects with the significantly increased anti-inflammatory cytokines in the serum of BD subjects as compared with CT, (t.test, p = 0.0001083). (T) GSVA score of the DEG from Zandi et al. in the anterior cingulate cortex of BD and CT subjects with the significantly decreased anti-inflammatory cytokines in the serum of BD subjects as compared with CT, (t.test, p = 0.04423). (correspondence of p value in graphs: *: p < 0.05, **: p < 0.01, *** p < 0.001, **** p < 0.0001).

